# Catestatin induces glycogenesis by stimulating phosphoinositide 3-kinase-AKT pathway

**DOI:** 10.1101/2020.10.31.363481

**Authors:** Gautam Bandyopadhyay, Kechun Tang, Nicholas J.G. Webster, Geert van den Bogaart, Sushil K. Mahata

## Abstract

**Aim:** Defects in hepatic glycogen synthesis contribute to postprandial hyperglycemia in type 2 diabetic (T2D) patients. Chromogranin A (CgA) peptide Catestatin (CST: hCgA_352-372_) has been shown to improve glucose tolerance in insulin-resistant mice. Here, we seek to determine whether CST also reduces hyperglycemia by increasing hepatic glycogen synthesis.

**Methods:** We determined liver glycogen, glucose-6-phosphate (G6P), uridine diphosphate glucose (UDPG), and glycogen synthase (GYS2) activities; plasma insulin, glucagon, norepinephrine (NE), and epinephrine (EPI) levels in fed and fasted liver of lean and obese mice as well as in CST knockout (CST-KO) mice after treatments with saline, CST, or insulin. We also determined glycogen synthesis and glycogenolysis in primary hepatocytes. In addition, we analyzed phosphorylation signals of Insulin receptor (IR), insulin receptor substrate-1 (IRS-1), phosphatidylinositol dependent kinase-1 (PDK-1), GYS2, glycogen synthase kinase-3β (GSK-3β), AKT (an enzyme in AKR mouse that produces Thymoma)/PKB (protein kinase B) and mTOR (mammalian/mechanistic target of rapamycin) by immunoblotting.

**Results:** CST stimulated glycogen accumulation in fed and fasted liver and in primary hepatocytes. CST reduced plasma NE and EPI levels, suggesting that CST promotes glycogenesis by inhibiting catecholamine-induced glycogenolysis. CST also directly stimulated glycogenesis and inhibited NE and EPI-induced glycogenolysis in hepatocytes. CST elevated the levels of UDPG and increased GYS2 activity, thus redirecting G6P to the glycogenic pathway. CST-KO mice had decreased liver glycogen that was restored by treatment with CST, reinforcing the crucial role of CST in hepatic glycogenesis. CST can improve insulin signals downstream of insulin receptor IR and IRS-1 by enhancing phospho-AKT signals through stimulation of PDK-1 and mTORC2 (mTOR complex 2) activities.

**Conclusions:** We conclude that CST directly promotes the glycogenic pathway and reduces plasma glucose levels in insulin-resistant mice by (i) reducing glucose production, (ii) increasing glycogen synthesis from UDPG, and (iii) reducing glycogenolysis. This is achieved by enhancing downstream insulin signaling.

## Introduction

Obesity, a major risk factor for type 2 diabetes (T2D) ^1^, results in the development of insulin resistance and post-prandial hyperglycemia ^2–4^. In the United States, >50% of the population have either diabetes or prediabetes ^5^. Obesity-induced T2D is thus emerging as a global health problem threatening to reach pandemic levels by 2030 ^6,7^. Chronic hyperglycemia causes glucotoxicity ^8,9^, which results in decreased secretion of insulin and increased insulin resistance ^8,10^. The liver plays a central role in maintenance of blood glucose homeostasis. In the postprandial state, the liver responds to an increase in portal vein blood glucose and insulin levels by absorbing glucose via glucose transporter 2 (GLUT2) and converting it to glycogen. The stimulation of net glycogen synthesis is a major, direct physiological function of postprandial insulin on hepatocytes ^11^. In the postabsorptive state, the liver produces glucose by glycogenolysis and gluconeogenesis ^12^. In the liver, glucose metabolism is determined largely by the glucose concentration in the portal vein (substrate supply) and is regulated by feed-forward activation mechanisms ^13,14^. Liver glycogen synthesis is impaired in T2D patients ^14,15^. Glycogen synthesis is also impaired in other organs, as for instance the reduced glucose uptake in skeletal muscle in T2D subjects is partly accounted for by reduced glycogen synthesis ^16^. Defects in the hepatic glycogen synthesis have been shown to contribute to postprandial hyperglycemia of patients with poorly controlled insulin-dependent diabetes mellitus ^15,17,18^ and non-insulin-dependent diabetic subjects ^19,20^. Therefore, targeting hepatic glucose storage or production is a potential therapeutic strategy for T2D.

Glucose homeostasis is controlled by several hormones, with insulin being the most important anabolic hormone and glucagon and catecholamines being the most catabolic hormones. Glucagon and catecholamines act by the following mechanisms: (i) inhibiting secretion of insulin via an α-adrenergic mechanism ^21^, (ii) stimulating glucagon secretion ^22^, (iii) stimulating hepatic glucose production (HGP) via both β- and α-adrenergic regulation of hepatic glycogenolysis and gluconeogenesis ^23–26^, (iv) mobilizing gluconeogenic substrates such as lactate, alanine, and glycerol from extrasplanchnic tissues to the liver ^27^, and (v) decreasing glucose clearance by direct inhibition of tissue glucose uptake ^28,29^. In addition, both glucagon and the catecholamines counteract insulin-induced hypoglycemia by stimulating glycogenolysis and gluconeogenesis and augmenting HGP ^30,31^.

Chromogranin A (CgA), a ~49 kDa proprotein, gives rise to peptides with opposing regulatory effects ^32–36^. While pancreastatin (PST: hCgA_250-301_) is an anti-insulin and proinflammatory peptide ^37,38^, catestatin (CST: CgA_352-372_) is a proinsulin and anti-inflammatory peptide ^39,40^ and is a risk factor for cardiovascular diseases in patients undergoing dialysis ^41^. The lack of all CgA peptides make the CgA knockout mice (CgA-KO) more sensitive to insulin despite being obese ^42^. In addition, opposite to wild-type (WT) mice, CgA-KO mice can maintain insulin sensitivity even after 4 months of high fat diet (HFD) ^43^. Lack of both CgA and Chromogranin B (CgB) also caused increased sensitivity to insulin with aging ^44^. In contrast, CST knockout (CST-KO) mice are obese and display insulin resistance and hyperinsulinemia even on normal chow diet (NCD) ^40^. We have recently shown that supplementation of CST-KO mice with CST not only restored insulin sensitivity but also normalized plasma insulin levels in CST-KO mice ^40^. In HFD-induced obese (DIO) mice, CST improved insulin sensitivity by attenuating inflammation, inhibiting infiltration of proinflammatory macrophages into the liver, and inhibiting gluconeogenesis ^40^. We have also shown that CST improves insulin sensitivity in DIO mice by attenuating HFD-induced endoplasmic reticulum stress ^45^. Another mechanism by which CST could potentially control hyperglycemia in insulin-resistant mice is by inducing synthesis of glycogen in the postabsorptive state.

Since CST inhibits both catecholamine secretion ^39,46–49^ and gluconeogenesis (by inhibiting expression of glucose 6-phosphatase gene (*G6pc*)) ^40^, we reasoned that glucose-6-phosphate (G6P), the substrate for *G6pc*, would accumulate after CST treatment, leading to stimulation of glycogenesis. To address this, we measured plasma insulin, glucagon, NE and EPI in CST-KO mice and DIO mice treated with CST and determined their effects on glycogen levels. This enabled us to determine whether the improved glucose tolerance that we reported earlier in DIO mice by CST ^40^, was due to CST’s interaction with pancreatic and adrenomedullary hormones leading to mobilization of G6P from the gluconeogenic to the glycogenic pathway. Therefore, in the present communication, we have tested the glycogenic effects of CST on genetically obese and insulin-resistant CST-KO mice as well as in insulin-resistant DIO mice and compared the effects of CST with insulin.

## Results

### CST induces hepatic glycogen synthesis in an insulin-independent pathway

One of the major pathways by which liver removes glucose after a meal is through insulin-induced conversion of glucose into glycogen (glycogenesis) ^10,11^. This is reflected by the presence of >3.5-fold more glycogen in fed liver compared to fasted liver (Fig. 1A), likely because insulin levels are reduced in fasted mice (Fig. 1B). Supplementation with insulin (30 minutes) restored glycogen level in fasted mice (Fig. 1A) Interestingly, like insulin, both acute (1 hour) and chronic (10-15 days) treatment with CST (2 μg/g body weight) raised glycogen levels in fasted mice (Fig. 1C). Moreover, CST treatment caused greater increase in glycogen levels compared to insulin even in fed mice (Fig. 1A&C). The combined treatment of insulin and CST elicited an additive effect on glycogen in fasted mice (Fig. 1D), suggesting that there may exist parallel pathways for CST and insulin actions. Ultrastructural changes of glycogen granules (GG) strengthen and support the biochemical findings of glycogen level (Fig. S1). GG granules in the liver were increased both in fed and fasted mice after CST treatment (Fig. S1 A-E). It was revealed that CST caused dose-dependent (10 ng to 25 μg/g body weight) increase in glycogen levels with an EC_50_ 0.013 μg (Fig. S1F). The half-life of CST in plasma in fed mice was approximately 6.7 hours (Fig. S1G).

**Figure 1.**
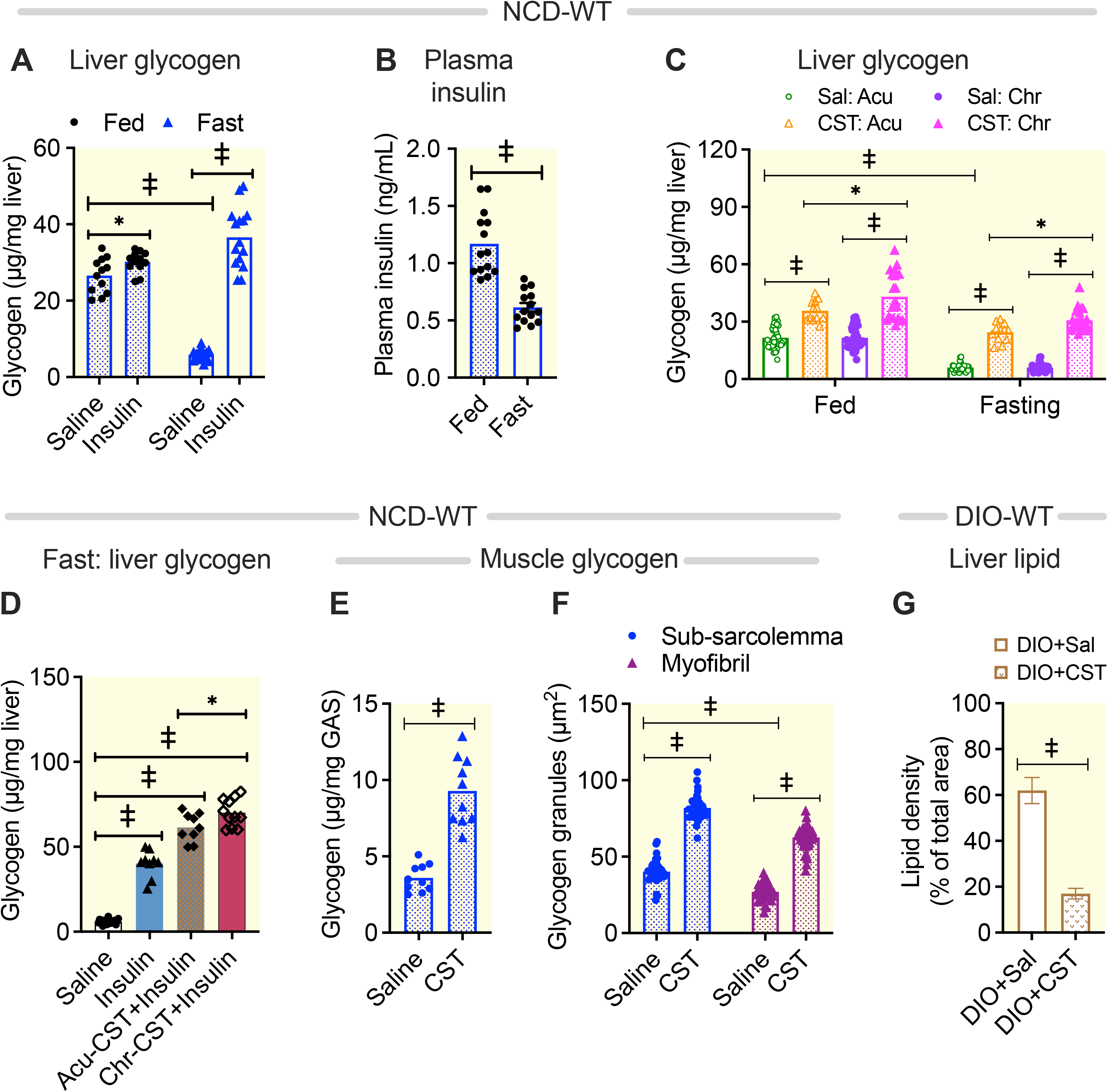
Effects of chronic CST and acute insulin treatments on hepatic glycogen contents in both fed and fasted NCD-WT mice. Mice were treated with CST (2 μg/g body weight/day for 15 days or 1 hour; intraperitoneal) followed by fasting (8 hr) and treatment with saline or insulin (0.4 mU/g body weight for 30 min; intraperitoneal) before harvesting tissues under deep anesthesia. (A) Liver glycogen content in fed (n=12) and fasted (n=15) NCD-WT mice. Two-way ANOVA: Interaction: p<0.001; Food: p<0.001; Treatment: p<0.001. (B) Plasma insulin levels in fed and fasted WT-NCD mice (n=14). Student’s *t* test. (C) Liver glycogen content in fed and fasted WT-NCD mice treated with saline (n=43), acute CST for 1 hour (n=12) or chronic CST for 15 days (n=43). Three-way ANOVA: Food: p<0.001; Acute vs Chronic: p<0.001; Saline vs CST: p<0.001; Food x Acute vs Chronic: ns; Food x Saline vs CST: p<0.05; Acute vs Chronic x Saline vs CST; p<0.001; Food x Acute vs Chronic x Saline vs CST: ns. (D) Liver glycogen content after saline (n=20), insulin for 30 min (n=9), acute CST plus insulin (n=9) or chronic CST for 15 days plus insulin for 30 min (n=11) treatment in fasted NCD-WT mice. One-way ANOVA. (E) Glycogen content in fed NCD-WT gastrocnemius muscle after saline or CST (2 μg/g body weight/day for 15 days; intraperitoneal) treatment (n=10). Student’s *t* test. (F) Morphometric analysis of 2-D TEM micrographs showing glycogen granules in saline and CST treated sub-sarcolemma and myofibril. Two-way ANOVA: Interaction: p<0.01; Treatment: p<0.001; Zone: p<0.001. (G) Morphometric assessment of lipid content in the TEM micrographs of steatotic liver of DIO mice after saline or chronic CST treatments. Student’s *t* test. *p<0.05; †, p<0.01; ‡, p<0.001.

### CST elevates muscle glycogen content

The bulk of postprandial glucose uptake takes place in the skeletal muscle ^50^. While ~80% of the glucose that enters muscle fibers in response to insulin is converted to glycogen ^51^, the rest is being oxidized to provide energy for muscle function ^16^. Being an insulin-sensitizing peptide ^40^, CST increased glycogen content in gastrocnemius muscle of NCD-fed WT mice (Fig. 1E). Electron microscopy revealed three distinct intracellular pools of glycogen ^52^: (i) subsarcolemmal glycogen, just beneath the sarcolemma, (ii) intermyofibrillar glycogen, located beneath the myofibrils (Fig. S2 A & B); and (iii) intramyofibrillar glycogen, in the myofibril, mainly near the z-line. Exercise has been reported to cause marked depletion of intramyofibrillar glycogen, underscoring glycogen’s key roles in muscle function ^53^. CST increased the number of glycogen granules both in the subsarcolemmal and intramyofibrillar region (Fig. 1F, S2A&B). As previously shown ^40^, CST decreased lipid density in the liver of DIO mice (Fig. 1G, S2E&F), thereby reducing steatosis.

### Contribution of catecholamines and glucagon

Glycogenolysis, controlled by the counterregulatory hormones such as glucagon, NE and EPI ^54^, helps the liver to maintain blood glucose concentrations during the early stage of fasting. We measured the effect of CST on these hormones. While plasma NE and EPI levels were reduced upon CST treatment in both fed and fasted state, no changes in glucagon levels were detected in the fed state (Fig. 3A). These findings suggest that one of the mechanisms by which CST restores and increases hepatic glycogen content in the early stage of fasting, could be by inhibiting catecholamine-induced glycogenolysis. Interestingly, acute insulin treatment (for 30 minutes) raised plasma NE and EPI levels only in the fasted state but not in the fed state (Fig. 3B&C). Exogenously administered insulin therefore seems to play a dual role in the fasting state to maintain glucose levels, stimulating gluconeogenesis by catecholamines but also promoting glycogenesis (Fig. 1A&D). Since in the fasted state, insulin not only increased liver glycogen content, but also increased plasma catecholamines, insulin mediated inhibition of glycogenolysis dominates over catecholamine induced glycogenolysis.

Since the glycogenic function of the liver is compromised in T2D ^19,20^, we tested the effects of CST on the hepatic glycogen content in insulin-resistant DIO mice. A schematic diagram shows the diet, peptide treatment and tissue harvesting (Fig. 3D). Chronic treatment of DIO mice with CST caused a >1.5-fold increase in liver glycogen content (Fig. 3E). When values in Figure 2E and F were compared, it appeared that in the fasted state, CST treatment increased liver glycogen content more than insulin in DIO mice. Thus, CST was more effective in inducing glycogenesis than insulin in DIO mice. Like in NCD mice, the effects of combined CST and insulin treatment in hepatic glycogen content were additive, indicating that these compounds both promote glycogenesis via divergent and synergistic pathways (Fig. 3F). We have previously shown that CST inhibits nicotine-induced secretion of NE from PC12 cells ^39,46,48^ as well as NE and EPI from mouse ^47^. Unlike in NCD mice, CST caused a decrease only in NE levels but did not change EPI and glucagon levels in fed DIO mice (Fig. S3). In fasted mice, the effects of CST on counterregulatory hormones were comparable between NCD and DIO mice (Fig. 2A&S3). Importantly, insulin-induced glycogen synthesis is reduced in DIO mice, but CST is equally effective in NCD or DIO mice. (Figs. 1C&3E).

**Figure 2.**
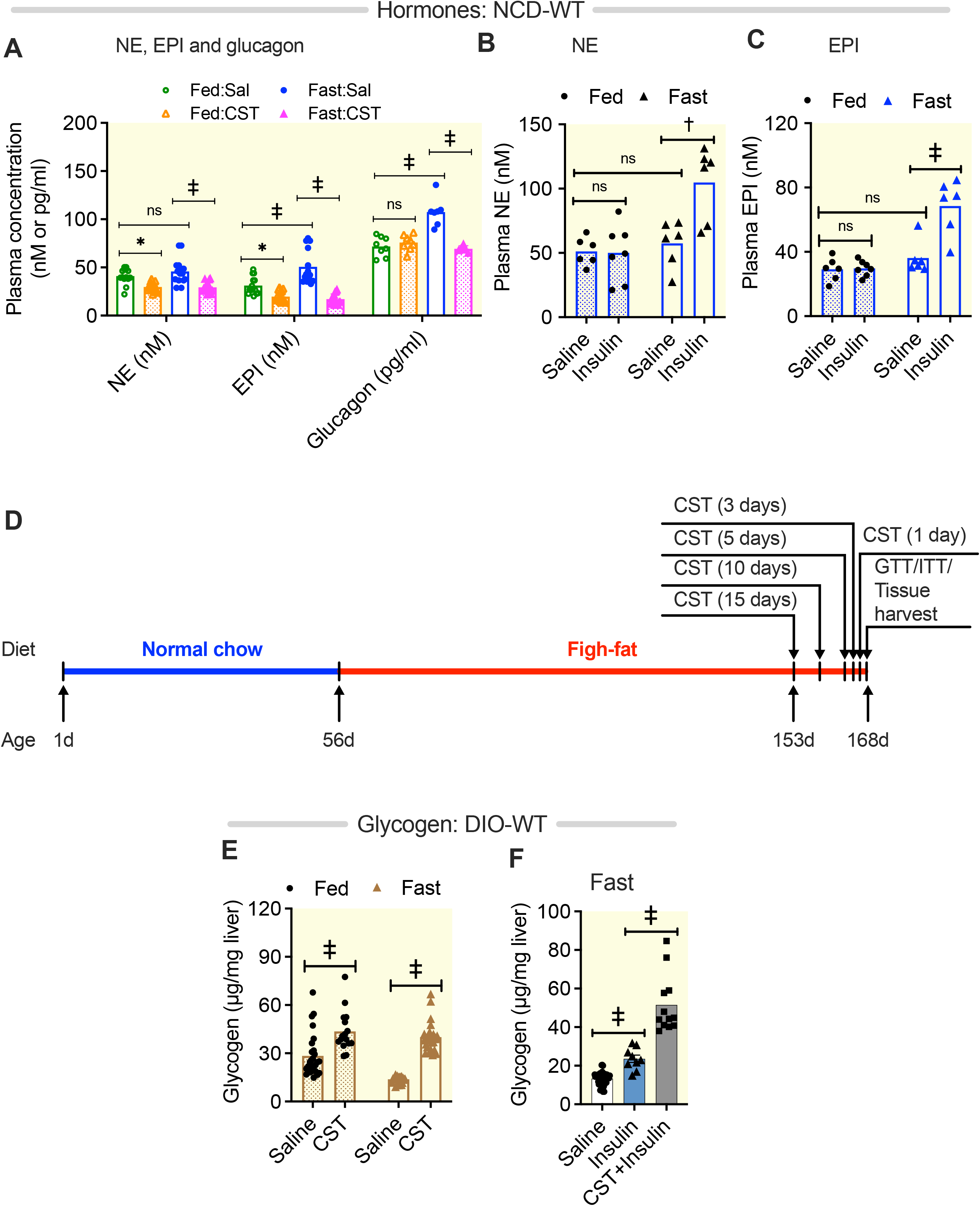
Effects of chronic CST and acute insulin treatments on plasma counterregulatory hormone levels in both fed and fasted NCD-WT mice. Fed or fasted NCD-WT mice were treated with CST (2 μg/g body weight/day for 15 days: intraperitoneal) before harvesting tissues under deep anesthesia after 24 hrs of last injection. (A) NE (n=16), EPI (n=16) and glucagon (n=6) levels in fed and fasted NCD-WT mice. Three-way ANOVA: Hormones: p<0.001; Fed vs Fast: p<0.001; Saline vs CST: p<0.001; Hormones x Fed vs Fast: p<0.05; Hormones x Saline vs CST: ns; Fed vs Fast x Saline vs CST: p<0.001; Hormones x Fed vs Fast x Saline vs CST: p<0.001. (B) NE levels in saline or insulin treated fed or fasted NCD-WT mice (n=7). Two-way ANOVA: Interaction: p<0.01; Treatment: p<0.05; Food: p<0.01. (C) EPI levels in saline or insulin treated fed or fasted NCD-WT mice (n=7). Two-way ANOVA: Interaction: p<0.01; Treatment: p<0.001; Food: p<0.001. (D) Schematic diagram showing age, diet, CST treatments, glucose and insulin tolerance tests, and tissue harvesting. (E) Liver glycogen content in fed and fasted DIO-WT mice after saline (n=27) or CST (n=16) treatments. Two-way ANOVA: Interaction: p<0.01; Treatment: p<0.001; Food: p<0.001. (F) Effects of saline (n=47), insulin alone (n=9; 30 min) or in combination with chronic CST (n=12) on glycogen content in fasted DIO-WT mice. One-way ANOVA: p<0.001. †, p<0.01; ‡, p<0.001; ns, not significant.

**Figure 3.**
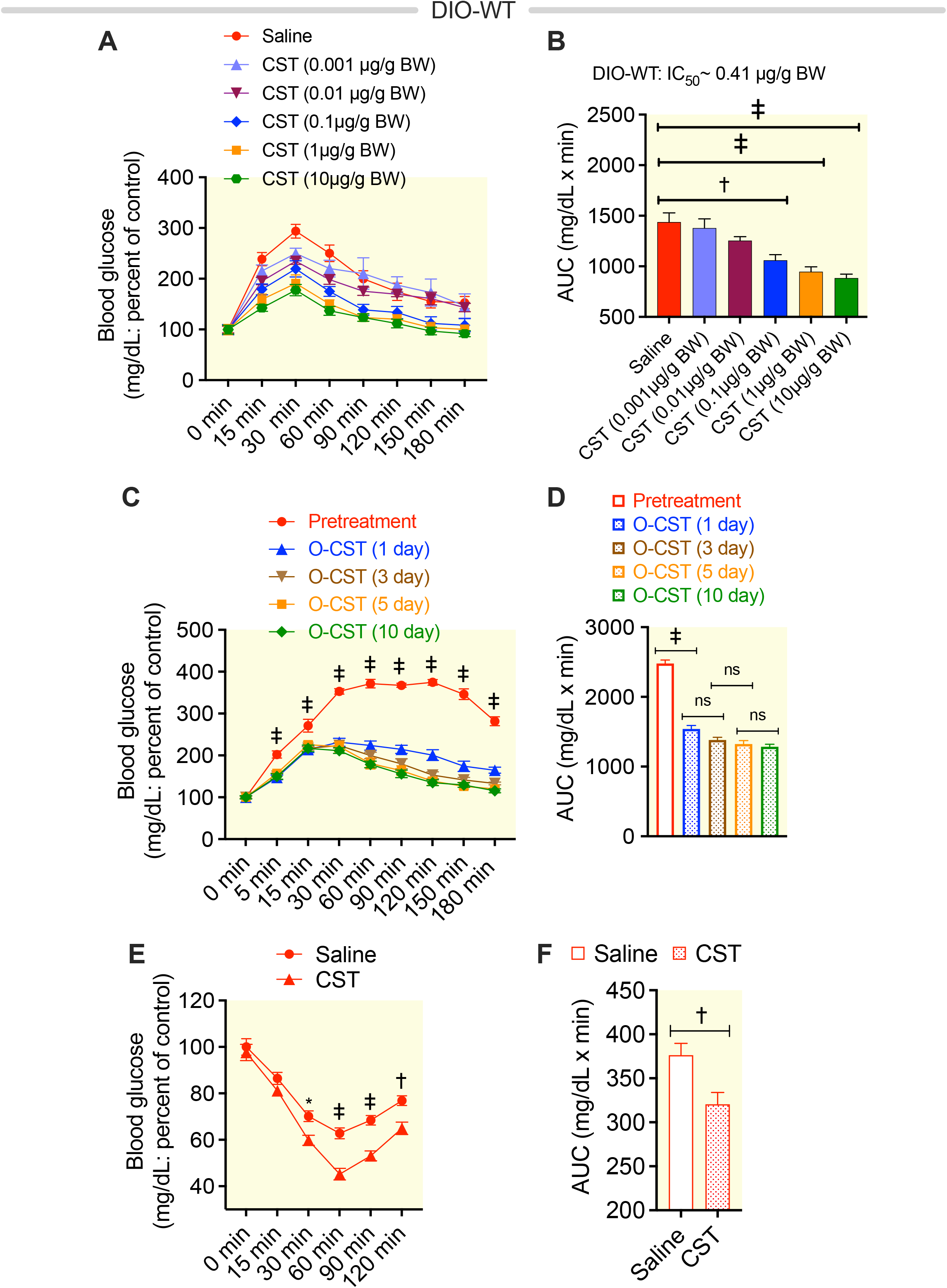
Effects of intraperitoneal or oral CST treatment on glucose tolerance in DIO mice without causing weight loss. (A&B) Oral glucose tolerance test (O-GTT): (A) Dose-dependent effects of oral CST on blood glucose levels during O-GTT (n=6). Two-way ANOVA: Interaction: p<0.05; Time: p<0.001; Treatment: p<0.001. (B) The area under the curve (AUC) of the OGTT was used to determine EC_50_ of CST action. One-way ANOVA: p<0.001. (C): Blood glucose levels during O-GTT after time-course of CST treatment (2 μg/g body weight) (n=7). Two-way ANOVA: Interaction: p<0.001; Time: p<0.001; Treatment: p<0.001. (D) The area under the curve (AUC) of the O-GTT. One-way ANOVA: p<0.001. (E) Intraperitoneal insulin tolerance test (ip-ITT): Blood glucose levels during ip-ITT after chronic CST treatment (n=12). Two-way ANOVA: Interaction: p<0.001; Time: p<0.001; Treatment: p<0.001. (F) The area under the curve (AUC) of the ip-ITT. Student’s *t* test. *p<0.05; †, p<0.01; ‡, p<0.001; ns, not significant.

### Dose dependent improvement of glucose tolerance by CST in DIO mice

We have recently shown that CST reduces inflammation in DIO mice, resulting in an improvement in insulin sensitivity ^40^. Although in our previous work we demonstrated improvement of glucose tolerance by CST ^40^, here we have conducted a dose-response study of CST (0.1-1.0 μg/g body weight) on glucose tolerance by intraperitoneal glucose tolerance test (ip-GTT) (Fig. 3A&B), showing that CST caused dose-dependent improvement in glucose tolerance (Fig. 3A&B). We have also conducted a time-course of oral CST action by oral-GTT (Fig. 3C&D) demonstrating that in DIO mice, revealing improved glucose tolerance by CST after 1 day of treatment (Fig. 3C&D). The absolute blood glucose values at 0 min are as follows: Pretreatment: 149.9±7.32; 1 day after CST: 142.7±8.836; 3 consecutive days of CST treatment: 138.6±8.16; 5 consecutive days of CST treatment: 134.0±11.66; 10 consecutive days of CST treatment: 128.9±11.9. Oral delivery of CST did not result in a significant reduction of the body weight (Fig. S4A). The intraperitoneal insulin tolerance test (ip-ITT) showed improvement in insulin tolerance in DIO mice by CST (Fig. 3F&G). The absolute blood glucose values at 0 min are as follows: saline: 155.5±10.62; CST: 121.2±9.63. Oral CST treatment also suppressed the expression of genes coding for enzymes involved in the gluconeogenesis: the G6Pase gene (*G6pc*) and phosphoenol-pyruvate kinase gene (*Pck1*) (Fig. S4 B & C), indicating that CST will limit *de novo* formation and hydrolysis of G6P. Together, these findings corroborated our previous work ^40^ by showing in further details that CST improves the GTT and ITT and inhibits the gluconeogenesis in DIO mice.

### Lower hepatic glycogen content in CST-KO compared to WT mice

To gain a better understanding of the role of CST in hepatic glucose homeostasis, we examined NCD-CST-KO mice, which are genetically obese and insulin-resistant ^40^. Hepatic glycogen content in fed NCD-CST-KO mice was lower than in WT mice (Fig. 4A), pointing to decreased insulin sensitivity. In the fed state, insulin failed to increase the hepatic glycogen content (Fig. 4B), supporting the presence of hepatic insulin resistance in CST-KO mice ^40^. In the fasted state, insulin caused only a ~1.4-fold increase in liver glycogen content in CST-KO mice (Fig. 4B). Chronic treatment of CST-KO mice with CST resulted in ~3-fold increase in hepatic glycogen content in fed mice and >4.2-fold increase in hepatic glycogen content in fasted mice (Fig. 4C). Insulin alone had no effect on accumulation of hepatic glycogen content in fasted NCD-CST-KO mice, reinforcing insulin resistance in CST-KO mice (Fig. 4D). Insulin in combination with CST caused significant increase in hepatic glycogen content in fasted NCD-CST-KO mice (Fig. 4D). Plasma insulin level in fasting mice were as follows: WT: 0.66±0.04 ng/ml; CST-KO: 1.59±0.12 ng/ml. Thus, fasting plasma insulin is CST-KO mice was >2.4-fold higher than WT mice, as shown in our previous work ^40^. Despite the presence of higher plasma insulin levels in the fasted state, fasted CST-KO mice showed a lower hepatic glycogen content than fasted WT mice (Fig. 4A). These effects of CST correlated with decreased levels of plasma counterregulatory hormones (Fig. 4F). Plasma glucagon levels in NCD-CST-KO mice were higher than DIO-WT mice (Figs. 4F& S3) but in contrast to DIO-WT mice, the glucagon levels were lower in CST-KO mice after CST treatment (Figs. 4F& S3).

**Figure 4.**
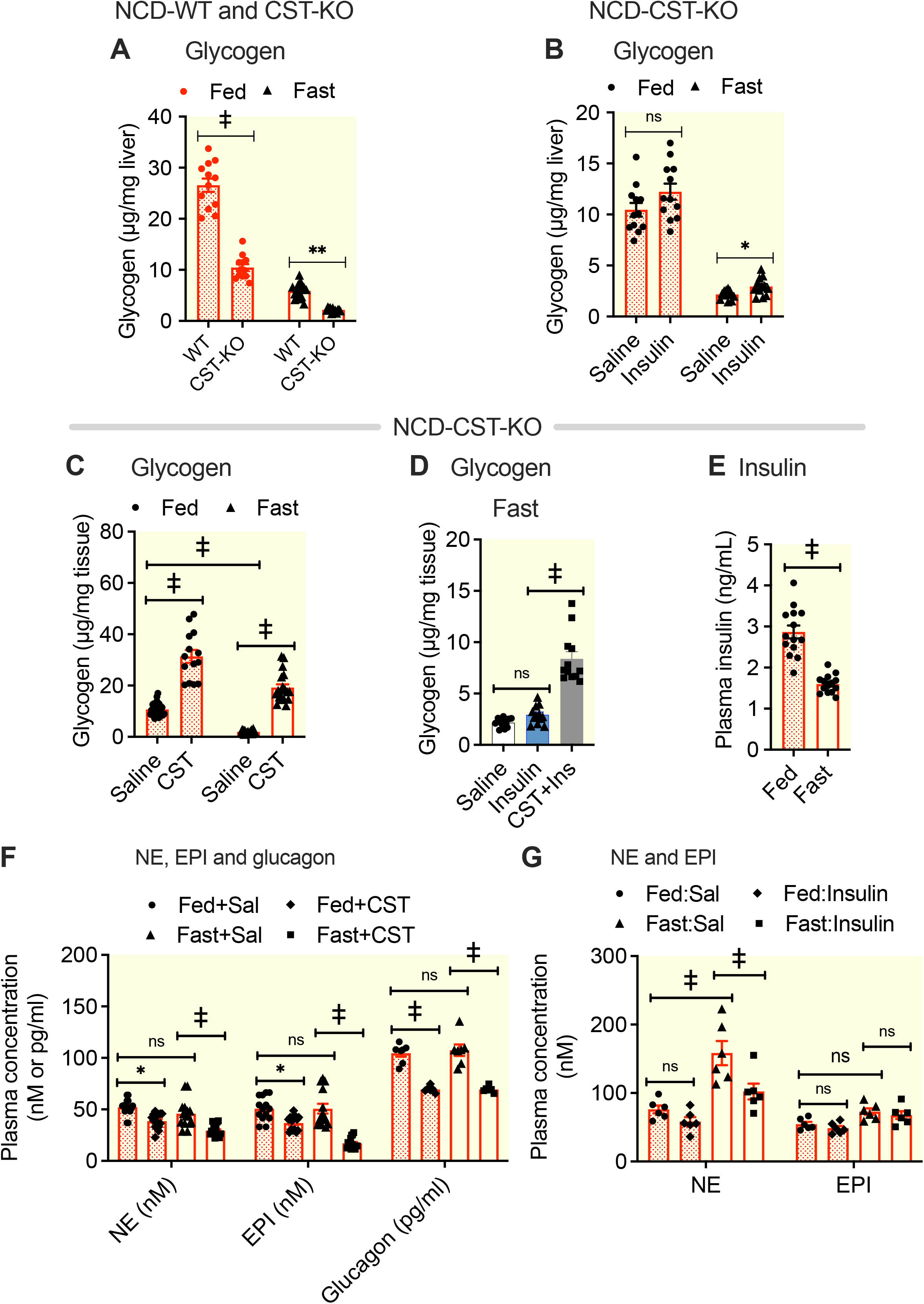
Effects of chronic CST and acute insulin treatments on hepatic glycogen content and plasma levels of counterregulatory hormones levels in NCD-CST-KO mice. NCD-WT and NCD-CST-KO mice were treated with CST (2 μg/g body weight/day for 15 days; intraperitoneal) followed by treatment with insulin (0.4 mU/g body weight for 30 minutes) before tissue harvesting. Tissues were harvested from fed or fasted (8 hr) mice under deep anesthesia. (A) Hepatic glycogen content in fed and fasted NCD-CST-KO mice compared to NCD-WT mice (n=12). Two-way ANOVA: Interaction: p<0.001; Treatment: p<0.001; Food: p<0.001. (B) Effects of insulin (30 min) on glycogen content in fed and fasted NCD-CST-KO mice (n=12). Two-way ANOVA: Interaction: ns; Time: Treatment: p<0.05; Food: p<0.001. (C) Liver glycogen content in fasted and fed NCD-CST-KO mice after saline (n=26) chronic CST treatment (n=14). Two-way ANOVA: Interaction: ns; Time: Treatment: p<0.05; Food: p<0.001. (D) Effects of insulin (30 min) alone or in combination with chronic CST on liver glycogen content in fasted CST-KO-NCD mice (n=12). One-way ANOVA: p<0.001. (E) Plasma insulin levels in fed or fasted NCD-CST-KO mice (n=14). Student’s *t* test. (F) NE (n=15), EPI (n=15), and glucagon (n=6) levels in fed or fasted NCD-CST-KO mice after chronic treatments with CST. Three-way ANOVA: Hormones: p<0.001; Fed vs Fast: p<0.01; Sal vs CST: p<0.001; Hormones x Fed vs Fast: ns; Hormones x Sal vs CST: p<0.001; Fed vs Fast x Sal vs CST: p<0.05; Hormones x Fed vs Fast x Sal vs CST: ns. (G) NE and EPI levels in saline or insulin (30 min) treated fed and fasted CST-KO-NCD mice (n=6). Three-way ANOVA: Catecholamines: p<0.001; Fed vs Fast: p<0.001; Sal vs Insulin: p<0.01; Catecholamines x Fed vs Fast: p<0.001; Catecholamines x Sal vs Insulin: p<0.05; Fed vs Fast x Sal vs CST: ns; Catecholamines x Fed vs Fast x Sal vs CST: ns. *p<0.05; ‡, p<0.001; ns, not significant.

Like in DIO-WT mice, CST treatment reduced plasma NE and EPI levels in both fed and fasted CST-KO mice (Fig. 4F). However, CST treatment of CST-KO mice also lowered glucagon levels (Fig. 4F), suggesting that CST might have aided glycogen accumulation by reducing glucagon-induced glycogenolysis. Acute insulin treatment only reduced NE (in contrast to WT mice) in fasted CST-KO mice but had no effects on NE or EPI in fed mice (Fig. 4G). Thus, CST-KO mice display lower hepatic glycogen content compared to WT mice, and this is elevated by CST supplementation.

### CST enhances glycogen synthesis and suppresses glycogenolysis

To confirm the *in vivo* studies and assess whether CST directly affects glycogenesis and glycogenolysis in hepatocytes, we measured glycogen synthesis from radiolabeled glucose and glycogenolysis from prelabeled stored glycogen in cultured primary hepatocytes. We found that CST and insulin alone caused a modest (>1.4-fold by CST; ~1.8-fold by insulin) increase in glucose incorporation into glycogen, while CST and insulin in combination resulted in a >2.6-fold increase in glycogen accumulation (Fig. 5A), consistent with a previous report ^12^. Analyzing the dose-dependent effects of CST on net glycogenesis, the EC_50_ turned out to be approximately 43.4 nM (Fig. 5B). We also found that combined treatments of counterregulatory hormones NE and EPI caused a ~2.3-fold increase in glucose release from prelabeled glycogen, which was inhibited by both CST and insulin (Fig. 5C), indicating that both CST and insulin can directly induce glycogen synthesis and inhibit catecholamine-induced glycogenolysis by hepatocytes.

**Figure 5.**
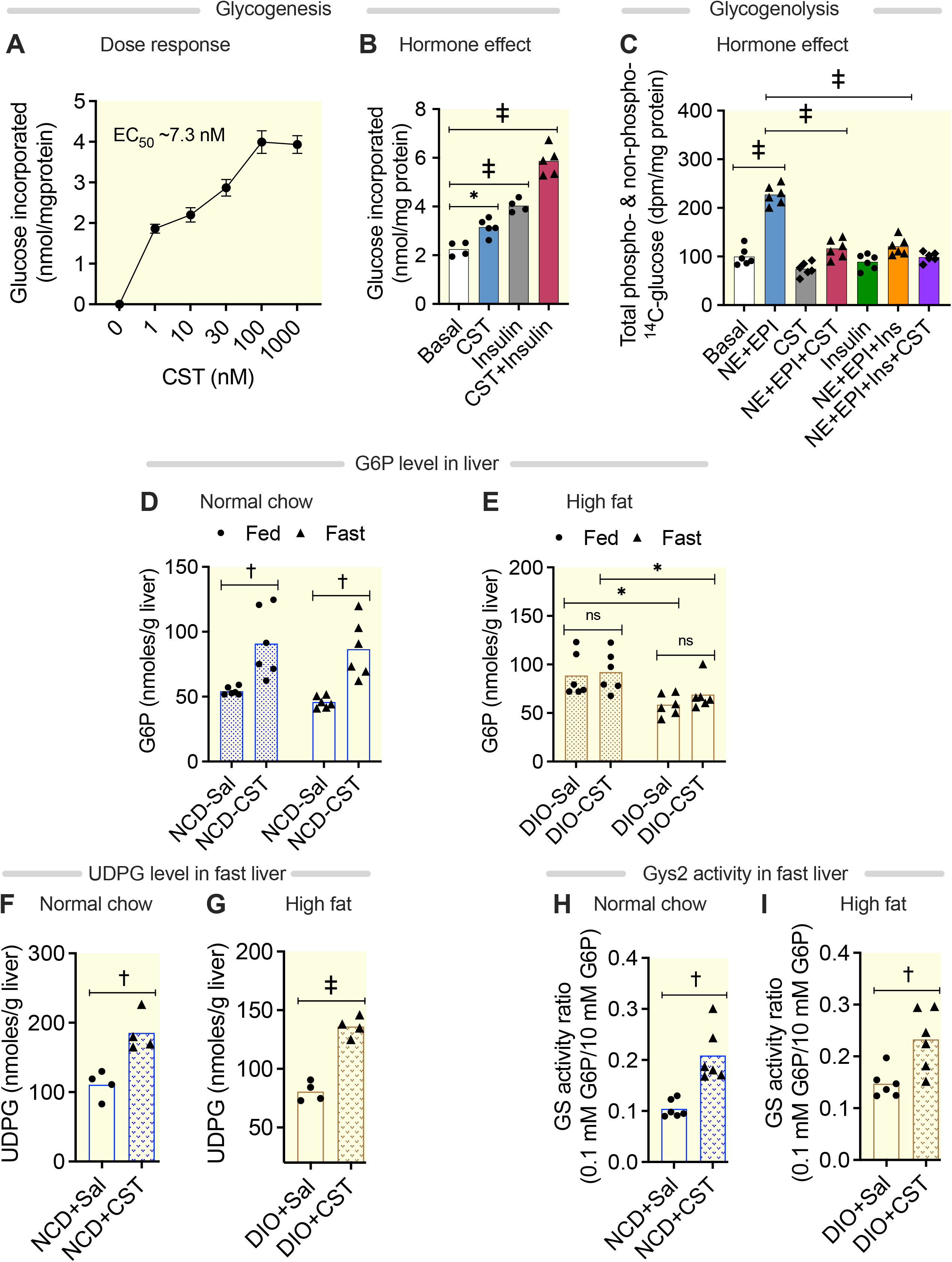
Regulation of glycogenesis and glycogenolysis in hepatocytes by CST, and regulation of the levels of G6P and UDPG as well as GYS2 activity in mouse liver by CST. (A) Glycogenesis from ^14^C-glucose in cultured primary lean hepatocytes (n=4-5). One-way ANOVA: p<0.001. (B) Dose-dependent effects of CST on glycogenesis. EC_50_ was calculated using GraphPad PRISM software. (C) Glycogenolysis from preloaded radiolabeled glycogen (n=6). One-way ANOVA: p<0.001. (D) Effects of CST on liver G6P levels in fed or fasted NCD-WT mice (n=6). Two-way ANOVA: Interaction: ns; Treatment: p<0.001; Food: ns. (E) Effects of CST on liver G6P levels in fed or fasted DIO-WT mice (n=6). Two-way ANOVA: Interaction: ns; Treatment: ns; Food: p<0.01. (F&G) Effects of CST on UDPG in liver of fasted NCD-WT or DIO-WT mice (n=4). Student’s *t* test. (H & I)) Effects of CST on GYS2 activities in liver of fasted NCD-WT and DIO-WT mice (n=6). Student’s *t* test. For GYS2 activity, the activation ratios in presence of low (0.1 mM) and high (10 mM) G6P are shown (n=6). Tissues were from the same experiments shown in Figures 1&2. *p<0.05; †, p<0.01; ‡, p<0.001, ns, not significant.

### CST increases G6P and UDPG levels

After entry of glucose into hepatocytes, glucose is immediately phosphorylated to G6P followed by conversion to glucose-1-phosphate (G1P) by phosphoglucomutase (PGM), and then to UDPG by UDP-glucose pyrophosphorylase to provide the substrate for glycogen synthesis. Glycogen synthase (GYS) or liver specific GYS2 catalyzes the transfer of glucosyl units from UDPG to glycogen by synthesis of α-1,4 bonds. A strong positive correlation exists between activation of GYS and intracellular G6P ^55,56^. Upon binding to GYS, G6P causes allosteric activation of the enzyme through a conformational rearrangement that simultaneously converts it into a better substrate for protein phosphatases ^57–59^. Dephosphorylation results in activation of GYS.

CST increased G6P levels in both fed (>1.68-fold) and fasted (>1.88-fold) NCD-WT mice (Fig. 5D). In contrast, CST did not increase G6P levels in both the fed and fasted states in DIO mice (Fig. 5E). However, the basal G6P levels in fed DIO mice seemed to be already higher than the basal levels in fed NCD mice, almost close to the levels of CST-treated NCD mice (Fig. 5E). Since UDPG levels were increased by CST in fasted NCD and DIO mice (Fig. 5 F & G), these results underscore the critical role of UDPG-pyrophosphorylase in CST action in DIO mice where G6P levels are not changed.

### CST augments GYS2 activity by inactivating GSK-3β through phosphorylation

GYS activity is controlled by covalent modification of the enzyme complex ^60^, allosteric activation by G6P ^58,61^, and enzymatic translocation by insulin ^62^. Therefore, we measured GYS2 activity by following incorporation of UDP-^14^C-glucose into glycogen in the presence or absence of saturating concentrations of G6P. CST stimulated GYS2 activity in fasted NCD-WT and DIO-WT mice (Fig. 5H &I). In addition to the allosteric activation of GYS2 by G6P, GYS2 is regulated by multi-site phosphorylation which causes inactivation. GYS2 is phosphorylated on nine serine residues by kinases such as glycogen synthase kinase (GSK)-3β ^63,64^. GSK-3β in turn is inactivated by phosphorylation of its Ser9 site by PKA, AKT and p90RSK and activated by autophosphorylation of Tyr216 ^65–68^. Phosphorylation of GYS2 results in its progressive inactivation and decreased sensitivity to allosteric activators ^69^. GYS2 dephosphorylation is regulated by hormones like insulin and is mediated by phosphatases like protein phosphatase 1 ^58^.

Immunoblotting revealed that in fasted NCD-WT and DIO-WT mice, CST and insulin alone increased GSK-3β Ser9 phosphorylation (Fig. 8A&B). The effects of CST in combination with insulin were additive in DIO-WT mice but not in NCD-WT mice (Fig. 8A&B). Since Ser641 is one of the GSK-3β target sites in GYS2, we also measured Ser641 phosphorylation by immunoblotting. In fasted NCD-WT mice, CST or insulin alone caused dephosphorylation of Ser641, while their combination caused a small additive effect (Fig. 8A&C). However, in DIO-WT mice, neither CST nor insulin reduced Ser641 phosphorylation, but the combination did (Fig. 8A&C). In summary, CST increases glycogen production in liver by increasing UDPG levels, increasing phosphorylation of GSK-3β, and dephosphorylation of GYS2.

### CST stimulates AKT signaling without affecting insulin-induced tyrosine phosphorylation of insulin receptor and insulin receptor substrate-1

#### (i) Activation of PI-3 kinase and PDK-1 dependent phosphorylation of Akt at threonine-308 residue

Insulin binding to the insulin receptor (IR) stimulates tyrosine phosphorylation (pY) of IR (through auto phosphorylation by IR tyrosine kinase) and insulin receptor substrate-1 (IRS-1) (by the IR tyrosine kinase). These are the key initial steps of the insulin signaling pathway ^70^. We tested whether CST has any effect on these early steps. As expected, Insulin stimulated tyrosine phosphorylation of IR and IRS-1 in cultured hepatocytes. However, the levels of phosphorylation were not significantly changed upon CST treatment (Fig. 6A-D), suggesting that the proximal part of the insulin signaling pathway is separate from the pathway for CST action. Since CST had no effects on the phosphorylation of IR and IRS1, we tested whether CST exerts any post-IR signaling. Interestingly, CST, like insulin, stimulated PI3-kinase activity (Fig. 6E). This enzyme primarily phosphorylates inositol lipids like phosphatidyl-inositol-phosphates such as PI-4,5 P (PIP2) at the 3 positions of the inositol moiety, to form PI-3,4,5 P (PIP3) which is a bioactive lipid. However, in this activity assay using anti-p85 immunoprecipitate (which pulls down catalytic subunit p110) as the source of the enzyme and phosphatidyl-inositol phosphate (PI), instead of PIP2, as the substrate, the product was PI3P (as shown in Fig. 6E). Thus, both insulin and CST treatment produced PI3P, and combined treatment with insulin and CST seems to have an additive effect. The endogenous products PIP2 and PIP3 binds to PH (Pleckstrin Homology) domain of AKT ^71^ as well as to mTORC2 ^72^. PIP2/PIP3 binding to Akt allows phosphorylation of AKT by phosphatidyl-inositol phosphate dependent kinase-1 (PDK-1) at threonine 308 residue ^71^. Downregulation of PDK-1 protein expression by si-RNA (Fig. 7A&B) led to the decreased pAKT (T308) signals (Fig. 7C&D). reenforcing the notion that the phosphorylation of Akt (T308), stimulated by insulin and CST, is PDK-1 dependent.

**Figure 6.**
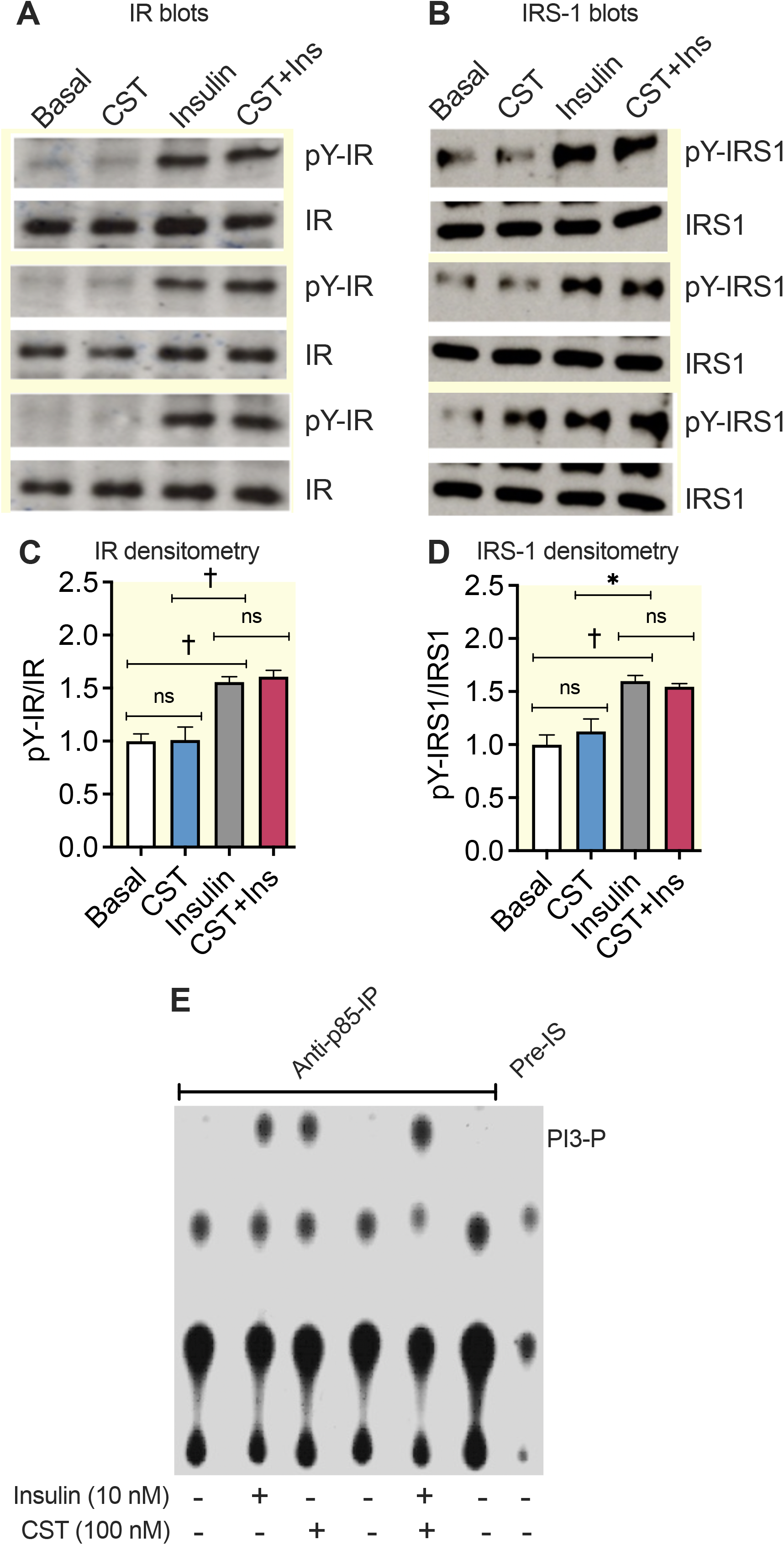
Unchanged tyrosine phosphorylation of IR and IRS-1 and increased PI3-kinase activity by CST in primary hepatocytes. (A) Immunoblots show tyrosine phosphorylation (pY) of IR in response to insulin and CST (n=3). (B) Immunoblots show tyrosine phosphorylation (pY) of IRS-1 in response to insulin and CST (n=3). (C) Corresponding density ratio of phospho-/total IR signals. One-way ANOVA: p<0.001. (D) Corresponding density ratio of phospho-/total IRS-1 signals. One-way ANOVA: p<0.01. (E) Autoradiograph of thin layer chromatography plate showing formation of PI3P from PI due to increased PI-3-kinase activity in the anti-p85-immunprecipitates after stimulation with insulin and CST. *p<0.05; †, p<0.01; ns, not significant.

**Figure 7.**
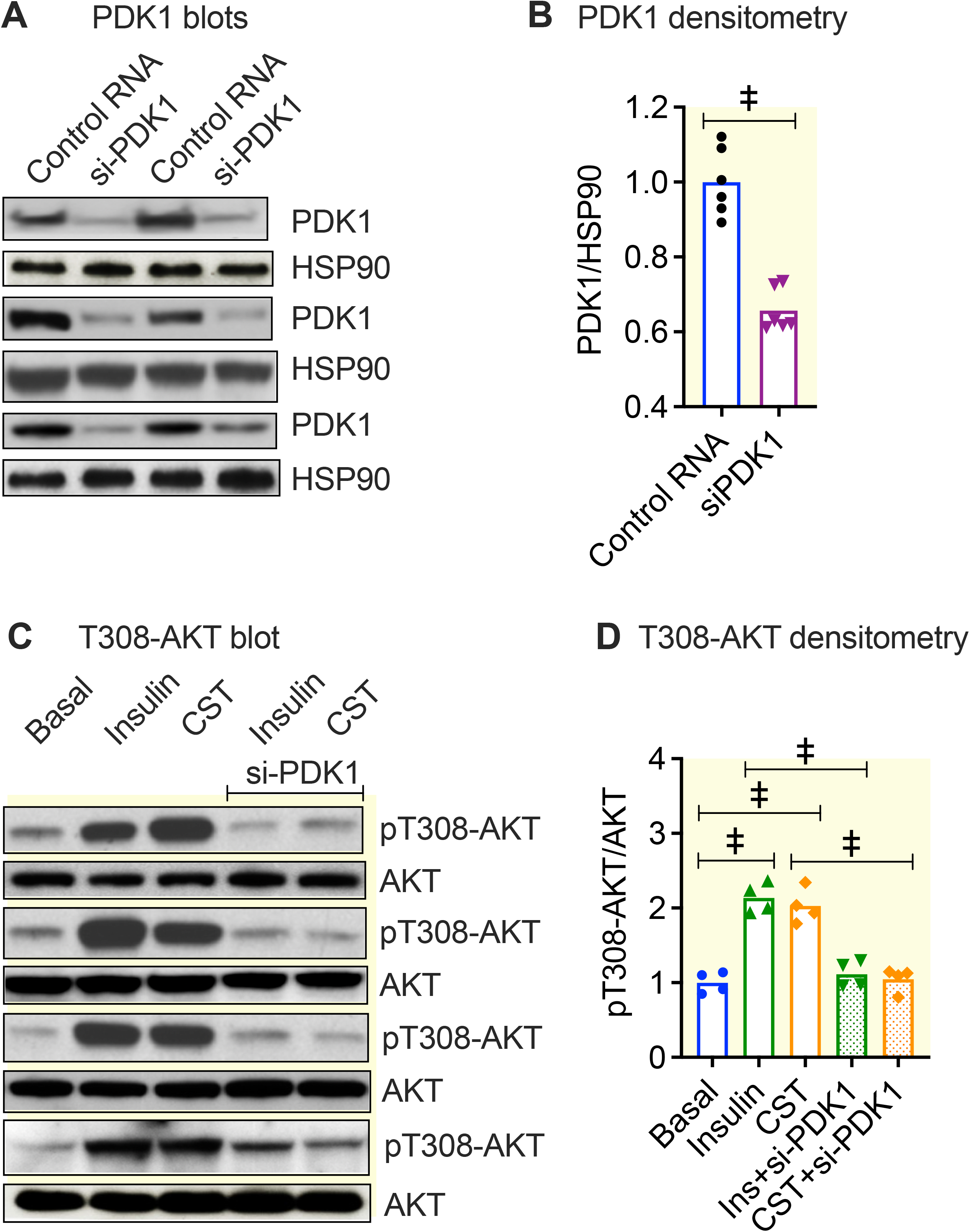
Increased phosphorylation of mTOR in the mTORC2 complex and AKT-S473 by insulin and CST in primary hepatocytes. (A) Immunoblots showing phosphorylation of mTOR at Ser2481 (n=3). (B) Corresponding density ratio of phospho-/total of mTOR signals. One-way ANOVA: p<0.001. (C) Immunoblots showing phosphorylation of AKT at Ser473 (n=3). (D) Corresponding density ratio of phospho-/total AKT signals are shown for AKT. One-way ANOVA: p<0.001. *p<0.05; †, p<0.01; ‡, p<0.001; ns, not significant.

#### (ii) Activation of mTORC2 and phosphorylation of AKT at the serine-473 residue

Another interesting aspect of the regulation of AKT phosphorylation is the existence of a positive feedback loop through which pT308-AKT phosphorylates SIN1 subunit of mTORC2 at the T86 residue enhancing mTORC2 kinase activity, which leads to phosphorylation of AKT at S473 (pS473-AKT) by mTORC2, thereby catalyzing full activation of AKT ^73^. Thus, mTORC2 is activated by both PIP3 as well as pT308-AKT. Furthermore, CST enhanced phosphorylation of mTOR at S2481 (pS2481) (Fig. 8A&B). This phosphorylation is known to increase functional integrity of the mTORC2 complex ^74^, causing increased AKT phosphorylation at S473 (Fig. 8C&D).

**Figure. 8.**
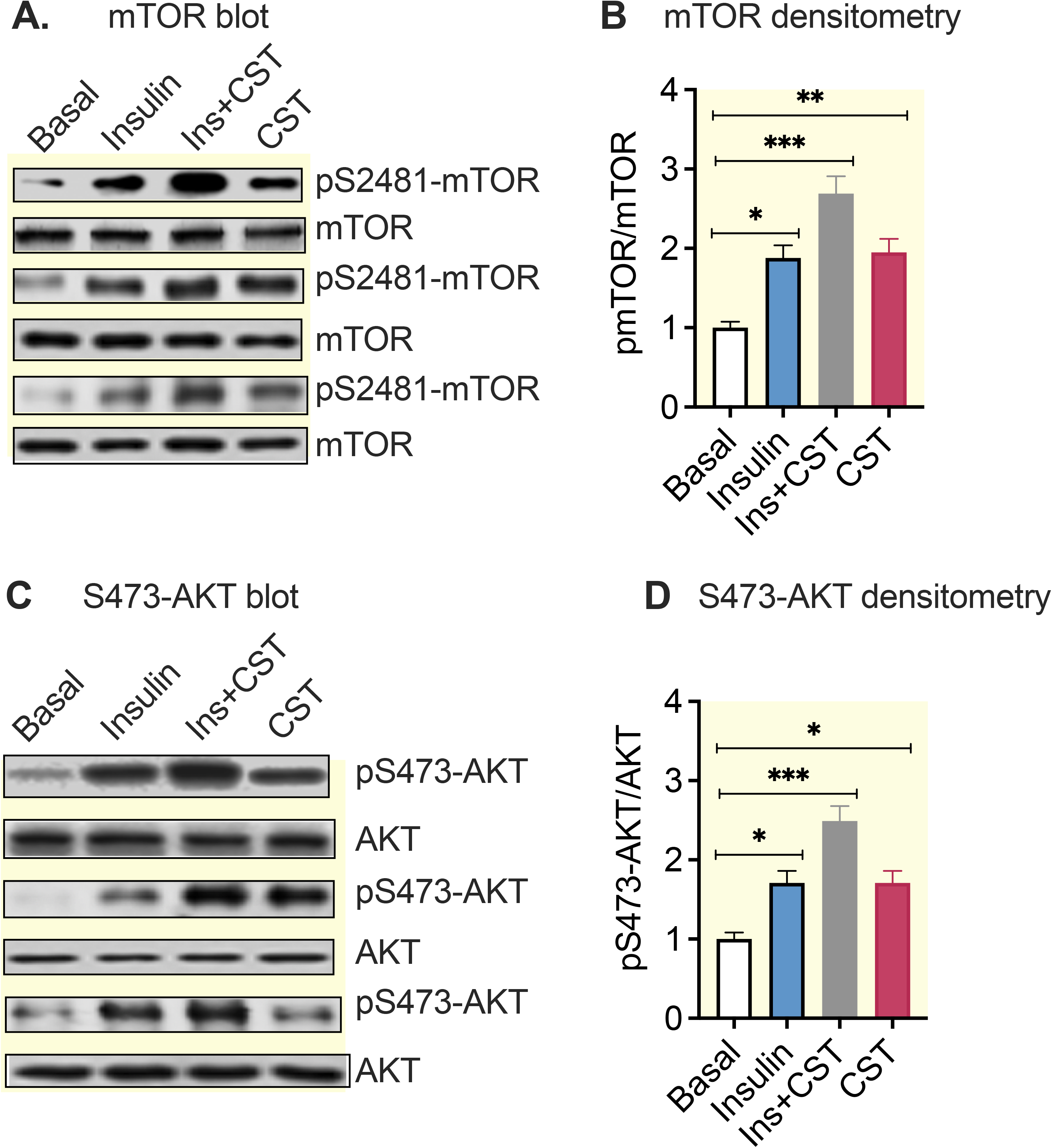
Stimulation of phosphorylation of AKT at T308 residue by insulin and CST in a PDK-1 dependent manner. (A) Immunoblots showing downregulation of PDK-1 protein expression by si-RNA. (B) The bar graphs showing signaling of the immunoblot. Student’s *t* test. (C) Immunoblots showing phospho-T308-AKT and total AKT signals after saline (Basal), insulin and CST treatments in presence and absence of si-RNA against PDK-1. (D) Bar graphs showing phospho-/total AKT signals. One-way ANOVA: p<0.001. ‡, p<0.001; ns, not significant.

Therefore, CST has a double acting role in AKT activation, one by providing PIP3 (via activation of PI3-kinase and PDK-1) and the other by strengthening the integrity of mTORC2 complex.

### CST augments GYS2 activity by inactivating GSK-3β through phosphorylation

Activation of Akt leads to phosphorylation of GSK-3β causing inactivation of GSK-3β which prevents subsequent phosphorylation and inactivation of GYS2. In addition to the allosteric activation, GYS2 is regulated by multi-site phosphorylation which causes inactivation. GYS2 is phosphorylated on nine serine residues by kinases such as glycogen synthase kinase (GSK)-3β ^63,64^. GSK-3β in turn is inactivated by phosphorylation of its Ser9 site by PKA, AKT and p90RSK and activated by autophosphorylation of Tyr216 ^65–68^. Phosphorylation of GYS2 results in its progressive inactivation and decreased sensitivity to allosteric activators ^69^. GYS2 dephosphorylation is regulated by hormones like insulin and is mediated by phosphatases like protein phosphatase 1 ^58^.

Immunoblotting revealed that in fasted NCD-WT and DIO-WT mice, CST, and insulin alone increased GSK-3β Ser9 phosphorylation (Fig. 9A&B). The effects of CST in combination with insulin were additive in DIO-WT mice but not in NCD-WT mice (Fig. 9A&B). Since Ser641 is one of the GSK-3β target sites in GYS2, we also measured Ser641 phosphorylation by immunoblotting. In fasted NCD-WT mice, CST or insulin alone caused dephosphorylation of Ser641, while their combination caused a small additive effect (Fig. 9A&C). However, in DIO-WT mice, neither CST nor insulin reduced Ser641 phosphorylation, but the combination did (Fig. 9A&C). In summary, CST increases glycogen production in liver by increasing UDPG levels, increasing phosphorylation of GSK-3β, and dephosphorylation of GYS2.

**Figure 9.**
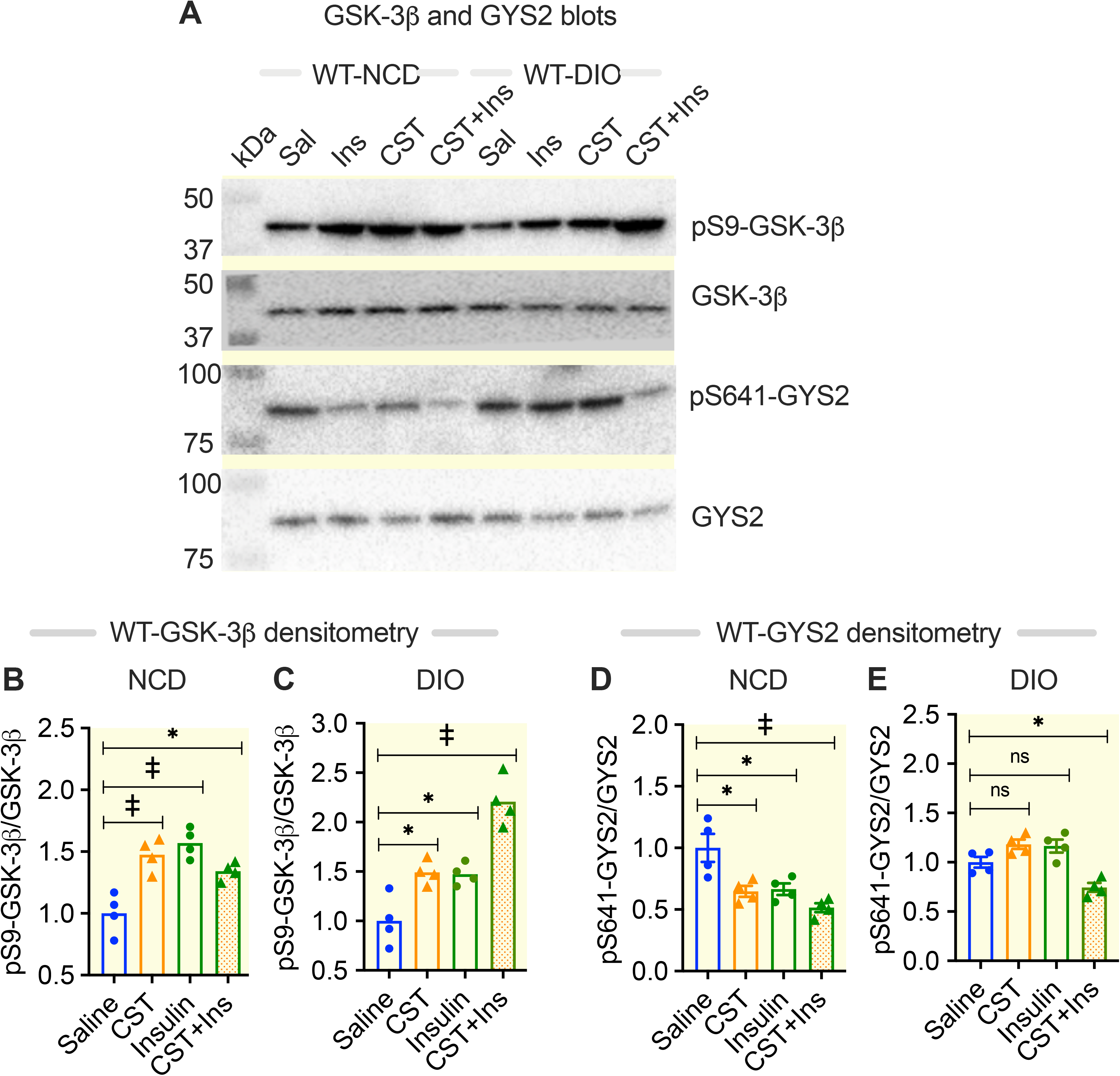
CST augments hepatic GYS2 activity (i.e., increased dephosphorylation) and decreases GSK-3β activity (i.e., increased phosphorylation) in fasted NCD-WT and DIO-WT mice. **(A)** Immunoblots showing total and phosphorylated signals for GSK-3β and GYS2 in NCD-WT and DIO-WT mice. **(B)** Densitometry values of GSK-3β in NCD-WT mice (n=4). One-way ANOVA: p<0.001. **(C)** Densitometry values of GSK-3β in DIO-WT mice (n=4). One-way ANOVA: p<0.001. **(D)** Densitometry values of GYS2 in NCD-WT mice (n=4). One-way ANOVA: p<0.01. **(E)** Densitometry values of GYS2 in DIO-WT mice (n=4). One-way ANOVA: p<0.001. Tissues were from the same experiments shown in Figures 1&2. *p<0.05; †, ‡, p<0.001; ns, not significant.

## Discussion

In the postprandial state, a major pathway that contributes to the removal of glucose from the portal vein by the liver is conversion of glucose into glycogen ^75,76^ and defects in hepatic glycogen synthesis contribute to postprandial hyperglycemia in patients with poorly controlled insulin-dependent diabetes mellitus ^15^.

The present study shows that CST induces hepatic glycogenesis in both the fed and fasted state in both lean (NCD) and obese (DIO) mice. CST caused an almost doubling of G6P levels in lean mice. In the liver of obese mice, basal G6P levels are already higher than in the liver of lean mice. Therefore, CST had no additional effects on G6P levels. In both lean and obese mice, GYS2 activity was stimulated by CST, resulting in the efficient conversion of G6P to glycogen. It is likely that the elevated G6P allosterically activates GYS to allow greater glycogen synthesis ^55,57,58^ after CST treatment. In fasted lean mice, CST or insulin alone induced >5-fold increase in hepatic glycogen content, while their combination caused an additive effect. This additive effect indicates that CST and insulin use divergent signaling pathways to induce hepatic glycogenesis in the postabsorptive state. Such an effect was limited in DIO mice because of insulin resistance. Nevertheless, CST also elevated glycogen content in the liver of DIO mice.

In CST-KO mice, supplemented CST reaches a steady state level of 4-5 nM concentration in plasma in about 15 hours. Physiologically, such a concentration of CST could sustain a significant increase in glycogen formation. Together with insulin, this concentration of CST can boost glycogen loading even in obese mice, thus reducing circulating glucose level, implicating that delivery of circulatory glucose from G6P will be reduced by suppressing expression of *G6pc* with consequent diversion towards glycogenic pathway.

Part of the CST effect *in vivo* could be indirect through changes in catecholamines. Indeed, CST decreased the levels of plasma catecholamines in the fed state, while insulin had no effect. However, CST did not change plasma glucagon levels. These findings indicate that CST restores and increases hepatic glycogen content by inhibiting catecholamine-induced glycogenolysis in the early stage of fasting. In contrast, insulin-induced hepatic glycogenesis is associated with increased plasma catecholamines in the fasted mice. The contribution of catecholamine-induced glycogenolysis to glucose production is well known ^23,77^. Suppression of catecholamine levels by CST can have dual indirect impacts as it could reduce both the gluconeogenic as well as the glycogenolytic effects of catecholamines ^77^.

Our results in hepatocytes showed that CST and insulin inhibit catecholamine-induced glycogenolysis. Although under stressed conditions (starvation) insulin elevates catecholamine levels, its inhibitory effects on glycogenolysis may still dominate allowing accumulation of glycogen. In cultured hepatocytes, both glucose and insulin suppress glycogenolysis by inhibiting glycogen phosphorylase a, in a process possibly involving protein phosphatase 1 ^78^. Protein phosphatase 1 inactivates glycogen phosphorylase via dephosphorylation of phosphorylase kinase (phosphorylated by PKA) as well as activates GYS2 by dephosphorylation ^79^. The net result of this is increased glycogen accumulation.

Studies in humans show a dichotomy of EPI and NE responses during the oral glucose tolerance test: decreased plasma EPI and increased plasma NE ^80^, where the authors did not find a correlation between changed NE and EPI with glucose tolerance in mice. CST deficiency in CST-KO mice creates glucose intolerance and insulin resistance even in NCD-fed CST-KO mice. As would be predicted, CST-KO mice stored less glycogen in the liver than WT mice. In the absence of CST, elevated pancreatic glucagon and higher plasma NE may contribute to the glucose intolerance which could be corrected by insulin or CST.

The glycogenic function of liver is compromised in both T2D subjects and DIO mice ^19,20^. CST increased the hepatic glycogen content in fed DIO mice and was more effective than insulin in inducing hepatic glycogenesis in fasted DIO mice. These findings indicate that the increased hepatic glycogenesis in both fed and fasted DIO mice by CST may be due to the increased insulin sensitivity that we have reported earlier ^40^. Besides the indirect effects through the catecholamines, CST seems to exert a direct effect on glycogenesis in hepatocytes. Both CST and insulin inhibit glycogenolysis in isolated hepatocytes but there is no additive effect, suggesting a common signaling pathway. In contrast, CST and insulin have an additive effect on glycogen accumulation that may indicate a divergence of CST and insulin effects at the level of glycogen synthesis. The fact that CST alone stimulates glycogen synthesis in insulin-resistant DIO mice and in isolated hepatocytes, supports a direct, insulin-independent regulatory role of CST in glycogenesis. CST treatment increased cellular G6P levels which is consistent with our previous finding that CST suppresses expression of the *G6pc* gene ^40^, thus reducing hydrolysis of G6P back to glucose. The present findings demonstrate that G6P is also directed by CST into the glycogenic pathway by stimulation of GYS and inhibition of glycogenolysis. Consistent with glucose intolerance in CST-KO mice ^40^, we found here the reduced ability in CST-KO mice to convert glucose to glycogen owing to lack of CST.

The nature of the relationship between insulin and CST-induced signaling pathways is incompletely understood. We show here that CST does not affect the proximal part of insulin signaling pathway, involving tyrosine phosphorylation of IR and IRS-1. On the other hand, under insulin resistant conditions (in DIO mice), insulin and CST cooperate to produce an additive effect on phosphorylation of GSK-3β and dephosphorylation of GYS2, suggesting that CST improves the weak insulin response in obese mice. How might this happen? Our results suggest a two-prong effect of CST: (i) stimulation of PI3-kinase-mediated production of PIP3 leading to increased phosphorylation of AKT at T308 in a PDK-1 dependent manner, and (ii) increased autophosphorylation of mTOR at S2481 leading to improved integrity of the mTORC2 complex which allows increased phosphorylation of AKT at Ser473 ^74,81^. A third mechanism could be the stimulation of a positive feedback loop, as suggested in the literature, where it was shown that activation of AKT (at T308) directly phosphorylate SIN1 subunit of mTORC2 complex and enhance phosphorylation of AKT at S473 ^73^. That may mean CST, like insulin, could be an operator of an activation loop between PDK-1 and mTORC2 leading to phosphorylation of AKT at two sites, T308 and S473. Thus, the canonical insulin pathway and this alternative pathway for CST action may explain the additive effects seen with a combination of insulin and CST. The pathways of actions of insulin and CST are illustrated in the Figure 10. The mTORC2 pathway and its effects on AKT signaling has been extensively discussed in the literature ^82^.

**Figure 10.**
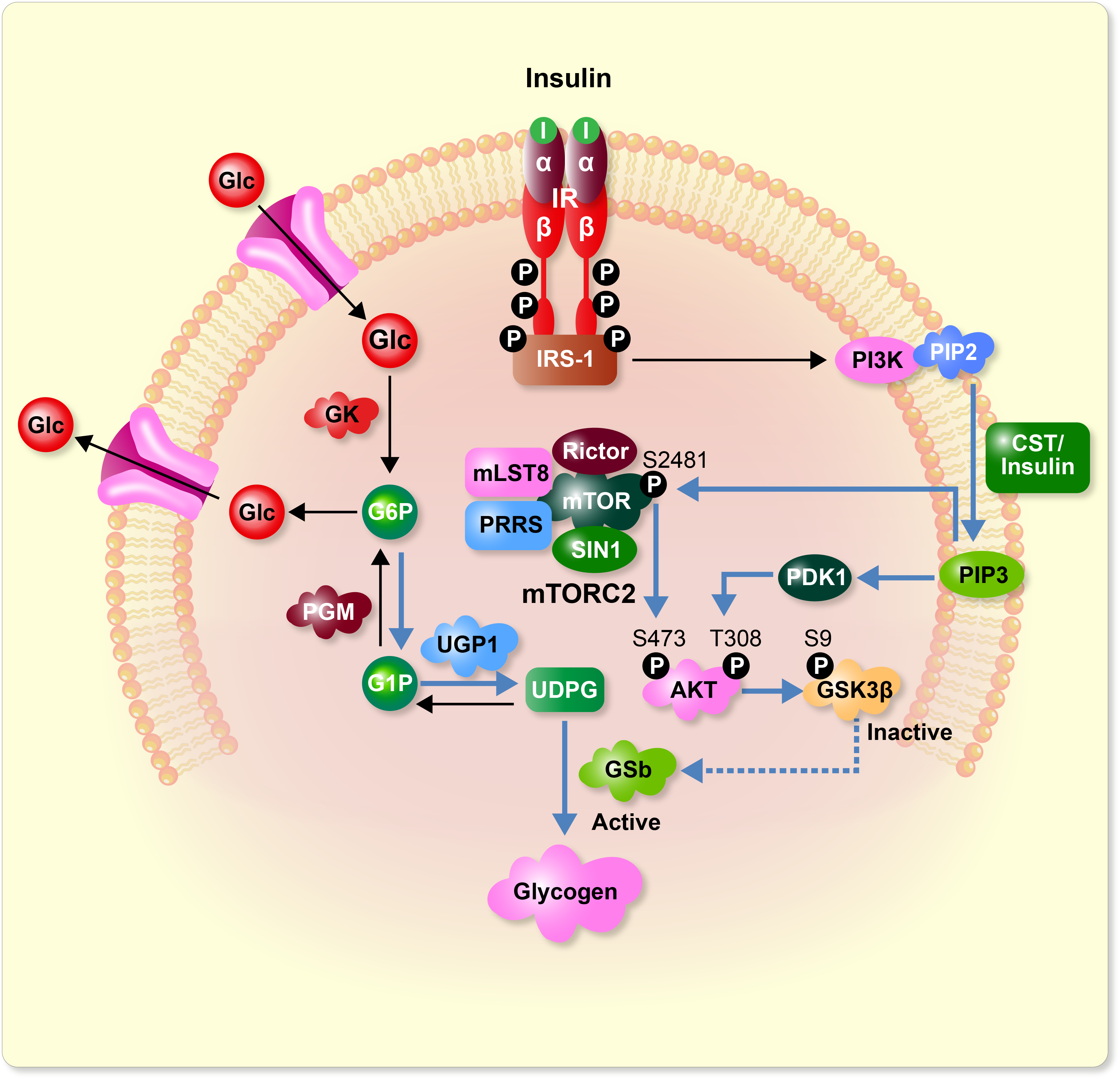
Schematic diagram showing insulin and CST mediated signaling pathways involved in the regulation of hepatic glucose and glycogen metabolism.

We have shown here three ways how CST causes glycogen accumulation. The first is to raise the substrate levels of glycogen synthesis (G6P and UDPG), one of which (G6P) is a known allosteric activator of GYS. The second is to enhance dephosphorylation of GYS by phosphorylating (and inactivating) GSK-3β. The third is to inhibit glycogenolysis. The insulin-independent stimulatory effect of CST on glycogenesis but not on glycogenolysis in hepatocytes, suggests that stimulation of glycogenesis by CST is the major pathway through which CST controls hepatic glucose production and improves glucose intolerance in obese mice. Given these characteristics, CST could be a new therapeutic peptide to treat both glucose intolerance and inflammation (in diabetes) as well as hypertension (due to its anti-adrenergic activity).

## Conclusion

CST improves glucose tolerance in obese and insulin-resistant mice by the following mechanisms: (i) reduced HGP from G6P, (ii) increased glycogen synthesis from G6P via formation of UDPG, (iii) phosphorylation of AKT at S473 and T308 by stimulating mTOR and PDK1, respectively, and (iv) reduced glycogen breakdown (due to suppression of plasma catecholamine levels). The net result is reduced free available glucose. These conclusions were verified in CST-KO mice and in hepatocyte cultures.

## Materials and Methods

### Animals, diets, and treatments

We used only male mice in this manuscript. Diet-induced obese (DIO) mice were created by feeding male WT mice (C57BL/6), starting at 8 weeks of age with HFD (Research diets D12492, 60% of calories from fat) for 112 days (16 weeks). Mice were kept in a 12:12 hour dark/light cycle; food and water were always available. Male CST knockout (CST-KO) mice with C57BL/6 background were also used. Both control WT mice as well as CST-KO mice were fed NCD (14% of calories from fat). Mice were injected intraperitoneally or orally with CST (2 μg/g body weight) for 1 hour (acute) on 168 days old mice (24 weeks) or 15 days (chronic) on day 153 days old mice. This dose of CST has been chosen from previous publications in rodents ^40,83–88^. Acute CST treatment (1 hour) was carried out in fasted mice 1 hour before sacrifice. A group of mice were fasted for 8 hours. Both fed and fasted mice were treated with saline or insulin (0.4 mU/g body weight) for 30 minutes before sacrificing for tissue collection that was done in the same way and at the same time of the day (8:00 to 9:30 AM). Tissues were rapidly frozen in liquid nitrogen and kept in −80°C freezer. Previously, we reported that CST treatment did not reduce body weight and did not change food intake in WT mice ^40^. In accordance with NIH animal care guidelines, all procedures and animals were housed and handled with the approval of The Institutional Animal Care and Utilization Committee.

### Glucose tolerance test (GTT) and insulin tolerance test (ITT)

For GTT, glucose (1 mg/g) was injected intraperitoneally (ip-GTT) or gavaged orally (O-GTT) (at time zero) after an 8-hour fast. Food was removed from the cages between 12:00 PM to 0:30 AM. Glucose and insulin were injected on fasted (8 hours) between 8:00 to 8:30 AM, respectively for GTT and ITT. Tail-vein glucose levels were measured using a glucometer at 0, 15, 30, 60, 90, and 120 minutes. For ITT, insulin (0.4 mU/g) was injected intraperitoneally, and blood glucose levels were measured using a glucometer at the indicated time points. Since plasma insulin level is low in fasting state, we injected insulin in fasted mice. GraphPad Prism software was used to determine the area under the curve (AUC) for each line curve.

### Hepatocyte isolation, culture, and assay for glycogen synthesis

Male mice (16-week-old) were fed NCD and used for perfusion of liver. Mice were perfused for 5 minutes with a calcium free buffer and followed by collagenase perfusion in a calcium containing buffer for another 5 minutes. Perfusion was carried out by inserting a catheter through inferior vena cava and passing buffer through a tube and allowing buffer to come out through portal vein which was cut for this purpose. The procedure was a modified version ^40^ of a published article ^89^. Livers, after collagenase digestion, were excised out, hepatocytes were squeezed out in a petri dish inside a culture hood, filtered through 100-micron nylon filter, centrifuged at 50 x g for 5 minutes, and pellets were collected. The suspensions of cell pellets were then passed through 30% isotonic percoll by centrifuging at 100 x g for 10 minutes. Pellets were washed in buffer and suspended in culture medium (Williams E) containing glutamax, 10% FBS, 10 nM dexamethasone and antibiotics. Hepatocytes were seeded on collagen I coated plates. Cells were cultured in Williams E medium and then depleted of glycogen store by incubating in Hepes Krebs Ringer Bicarbonate buffer for 8 hours followed by switching to serum-free Williams E medium (25 mM glucose) with or without the presence of insulin (10 nM), CST (100 nM) or insulin+CST, incubated for 24 hours in presence of ^14^C-glucose (10 μCi/mL, 5 mM). At the end of incubation, cultures were washed, and cells were dissolved in 30% KOH, boiled for 20 minutes, glycogen was precipitated by 60% ethanol (in presence of 5 μg glycogen as carrier), pellets washed with 70% ethanol, dried free of traces of ethanol, dissolved in water and counted.

### Glycogen and enzyme analysis in liver

Fresh and frozen liver (25-30 mg) and muscle (90-100 mg) tissues from fed or fasted mice (collected between 8:00 to 9:30 AM under deep anesthesia) and glycogen was extracted by boiling with 30% KOH solution as described previously ^90^. Extracted glycogen was precipitated in cold by 66% ethanol and washed with 70% ethanol. After drying to remove traces of ethanol, the pellets were dissolved in water and then subjected to colorimetric determination of glycogen using Anthrone reagent (0.05% anthrone and 1% thiourea) in concentrated H_2_SO_4_.

A group of lean and obese mice were fasted for 8 hours (12:00 PM to 0:30 AM) and treated with saline or insulin for 15 minutes. Mice were sacrificed (8:00 to 8:30 AM) and livers were subjected to analysis of GYS2 activity and immunoblotting. GYS2 activity was measured by incorporation of UDP-^14^C-glucose into glycogen in presence of 10 mM and 0.1 mM G6P, an activator, and ^14^C-glycogen formed was spotted on GF/A filter paper, washed with 70% ethanol, dried and counted for radioactivity. Counts were subsequently normalized against protein. Results are presented as activation ratio (0.1 mM G6P/10 mM G6P).

### Measurement of catecholamines

Blood was drawn from the heart between 8:00 – 9:30 AM under deep isoflurane-induced anesthesia (1-2 min) and kept in potassium-EDTA tubes. Plasma catecholamines were measured by ACQUITY UPLC H-Class System fitted with Atlantis dC18 reversed-phase column (100A, 3 μm, 2.1 mm x 150 mm) and connected to an electrochemical detector (ECD model 2465) (Waters Corp, Milford, MA) as described previously ^88^. The mobile phase (isocratic: 0.3 mL/min) consisted of phosphate-citrate buffer and acetonitrile at 95:5 (vol/vol). For determination of plasma catecholamines, DHBA (2 ng) was added to 150 μl plasma and adsorbed with ~15 mg of activated aluminum oxide for 10 min in a rotating shaker. After washing with 1 mL water adsorbed catecholamines were eluted with 100 μl of 0.1N HCl. 10 μl of the eluate was injected into UPLC where the ECD range was set at 500 pA. The data were analyzed using Empower software 3 (Waters Corp, Milford, MA). Catecholamine levels were normalized with the recovery of the internal standard. Plasma catecholamines are expressed as nM.

### Analysis of glycogenolysis in hepatocytes

Radiolabeled glycogen was synthesized in hepatocytes from ^14^C-glucose (10 μCi/mL) as described above in presence of insulin only (no CST). Cultures were washed (Krebs-Ringer bicarbonate buffer) to remove external radioactivity and then exposed to buffer (basal), NE (10 μM), EPI (10 μM), insulin (100 nM) and CST (100 nM) alone and in various combinations and incubated for 60 minutes. Incubation media were collected for radioactive counting. Cells were washed with cold buffer followed by cell lysis. An aliquot of each lysate was saved for protein assay and the rest was treated with 0.2 M barium hydroxide and 5% zinc sulphate (to precipitate protein) and centrifuged at 10,000 x g for 10 minutes. Supernatants were passed through 100 mg mixed bed ion-exchange resins (both cationic and anionic). Resins were washed twice with water and counted for bound radioactivity (glucose phosphates liberated from hydrolysis of radiolabeled glycogen will remain bound to the resins). For each incubation, total radioactivity was determined by combining the incubation medium, resin flow through and the radioactivity bound to the resin. The total radioactivity was normalized with protein, and it represented ^14^C-glucose (phosphorylated and non-phosphorylated) released from ^14^C-glycogen due to glycogenolysis.

### Measurement of G6P and UDPG

G6P levels were measured in liver tissues by a commercial kit (Sigma-Aldrich, St. Louis, MO) and UDPG levels were measured by an UPLC-UV detection method in the UCSD core facility.

### Assay for PI-3-kinase activity

CST (100 nM for 1hour), insulin (10 nM for 10 minutes) and untreated hepatocytes were lysed, and proteins were subjected to immunoprecipitation with anti-p85 antibody. After extensive washing, immunoprecipitates were analyzed for kinase activity in presence of phosphatidylinositol (PI) and γ-^32^P-ATP (PerkinElmer, Santa Clara, CA) followed a published method ^91^. Extracted radiolabeled lipids, mixed with reference lipids, were analyzed by TLC using a developing solvent, CHCl_3_–MeOH–H_2_O– NH_4_OH (25%), 45:35:8.5:1.5 (v/v). Reference spots were identified by exposing to iodine vapor in a closed chamber. TLC plates were exposed to X-ray film for 72 hours at −80°C. Reference spots for PI3P were counted for radioactivity.

#### Real Time PCR

Total RNA from liver samples was isolated using RNeasy Mini Kit and reverse-transcribed using qScript cDNA synthesis kit. cDNA samples were amplified using PERFECTA SYBR FASTMIX L-ROX 1250 and analyzed on an Applied Biosystems 7500 Fast Real-Time PCR system. All PCRs were normalized to *Rplp0*, and relative expression levels were determined by the ΔΔ*C_t_* method.

### Immunoblotting

Homogenization of livers were made in a lysis buffer supplemented with phosphatase and protease inhibitors. Homogenates were subjected to 10% SDS-PAGE and immunoblotted. The following primary antibodies were obtained from Cell Signaling Technology (Danvers, MA): GSK-3β (Catalog #9315) and pSer9-pGSK-3β (Catalog #5558), mouse monoclonal phospho-tyrosine 4G10 (Catalog # 96215), rabbit monoclonal GYS (Catalog #3886), and rabbit monoclonal, pSer641/640-GS (Catalog #3891). According to Cell Signaling Technology, this phospho-GYS antibody (Catalog #891) works to detect both muscle (pSer640) and liver (pSer641) signals. The antibodies against insulin receptor b (Catalog # sc-373975) and IRS-1 (Catalog # sc-8038) were purchased from Santa Cruz Biotechnology (Dallas, TX). Primary antibodies against PDK1, Heat shock protein-90 (HSP-900, p85 (a PI-3-kinase subunit), pT308-AKT, pS473-AKT were purchased from Cell Signaling Technology.

### Downregulation of PDK-1 by siRNA

Four siRNAs targeting mouse PDK-1 and a non-targeting siRNA pool were purchased from Dharmacon (Lafayette, CO). Hepatocytes were transfected with targeting and non-targeting siRNAs using Lipofectamine RNAiMax reagent (ThermoFisher Scientific) as the transfection reagent following manufacturer’s protocol. Transfection was continued for 48 hours before cells were homogenized. CST treatment (100 nM) was for 1 hour and insulin (10 nM) was for 10 minutes before cell homogenization.

### Statistical analysis

Statistics were performed with PRISM 8 (version 8.4.3) software (San Diego, CA). Normality was assessed by either D’Augustino-Pearson omnibus normality test or Shapio-Wilk normality test. After passing the normality test, data were analyzed using either unpaired two-tailed Student’s *t*-test for comparison of two groups or one-way or two-way analysis of variance (ANOVA) for comparison of more than two groups followed by Tukey’s *post hoc* test if appropriate. All data are presented as mean ± SEM and significance were assumed when p<0.05.

## Supporting information

Supplemental Figures_BioRxiv_110421

## Author contributions

GB researched the data, contributed to the design and discussion, and wrote the manuscript. KT generated the data, contributed to the design and discussion. NJW and GvdB edited the MS and provided intellectual inputs. SKM conceived the idea, generated ultrastructural data, supervised the studies, analyzed the data and made the graphics, wrote the MS, and provided financial support. GB and SKM are the guarantor of this study.

## Acknowledgments

This work was supported by grants from the Veterans Affairs (I01 BX003934 to SKM).

## Availability of data and material

The datasets generated during this study are available from the corresponding author on reasonable request.

## References

1. Martin BC, Warram JH, Krolewski AS, Bergman RN, Soeldner JS, Kahn CR. Role of glucose and insulin resistance in development of type 2 diabetes mellitus: results of a 25-year follow-up study. Lancet. 1992;340(8825):925–929.

2. Kahn SE, Hull RL, Utzschneider KM. Mechanisms linking obesity to insulin resistance and type 2 diabetes. Nature. 2006;444(7121):840–846.

3. Shoelson SE, Herrero L, Naaz A. Obesity, inflammation, and insulin resistance. Gastroenterology. 2007;132(6):2169–2180.

4. DeFronzo RA, Ferrannini E, Groop L, et al. Type 2 diabetes mellitus. Nat Rev Dis Primers. 2015;1:15019.

5. Menke A, Casagrande S, Geiss L, Cowie CC. Prevalence of and Trends in Diabetes Among Adults in the United States, 1988-2012. JAMA. 2015;314(10):1021–1029.

6. Zimmet P, Alberti KG, Shaw J. Global and societal implications of the diabetes epidemic. Nature. 2001;414(6865):782–787.

7. Wild S, Roglic G, Green A, Sicree R, King H. Global prevalence of diabetes: estimates for the year 2000 and projections for 2030. Diabetes Care. 2004;27(5):1047–1053.

8. Kawahito S, Kitahata H, Oshita S. Problems associated with glucose toxicity: role of hyperglycemia-induced oxidative stress. World J Gastroenterol. 2009;15(33):4137–4142.

9. Campos C. Chronic hyperglycemia and glucose toxicity: pathology and clinical sequelae. Postgrad Med. 2012;124(6):90–97.

10. Dubois M, Vacher P, Roger B, et al. Glucotoxicity inhibits late steps of insulin exocytosis. Endocrinology. 2007;148(4):1605–1614.

11. Roden M, Perseghin G, Petersen KF, et al. The roles of insulin and glucagon in the regulation of hepatic glycogen synthesis and turnover in humans. J Clin Invest. 1996;97(3):642–648.

12. Agius L. Physiological control of liver glycogen metabolism: lessons from novel glycogen phosphorylase inhibitors. Mini Rev Med Chem. 2010;10(12):1175–1187.

13. Cherrington AD. Banting Lecture 1997. Control of glucose uptake and release by the liver in vivo. Diabetes. 1999;48(5):1198–1214.

14. Moore MC, Coate KC, Winnick JJ, An Z, Cherrington AD. Regulation of hepatic glucose uptake and storage in vivo. Adv Nutr. 2012;3(3):286–294.

15. Hwang JH, Perseghin G, Rothman DL, et al. Impaired net hepatic glycogen synthesis in insulin-dependent diabetic subjects during mixed meal ingestion. A 13C nuclear magnetic resonance spectroscopy study. J Clin Invest. 1995;95(2):783–787.

16. Shulman GI, Rothman DL, Jue T, Stein P, DeFronzo RA, Shulman RG. Quantitation of muscle glycogen synthesis in normal subjects and subjects with non-insulin-dependent diabetes by 13C nuclear magnetic resonance spectroscopy. N Engl J Med. 1990;322(4):223–228.

17. Cline GW, Rothman DL, Magnusson I, Katz LD, Shulman GI. 13C-nuclear magnetic resonance spectroscopy studies of hepatic glucose metabolism in normal subjects and subjects with insulin-dependent diabetes mellitus. J Clin Invest. 1994;94(6):2369–2376.

18. Shulman GI. Cellular mechanisms of insulin resistance. J Clin Invest. 2000;106(2):171–176.

19. DeFronzo RA, Simonson D, Ferrannini E. Hepatic and peripheral insulin resistance: a common feature of type 2 (non-insulin-dependent) and type 1 (insulin-dependent) diabetes mellitus. Diabetologia. 1982;23(4):313–319.

20. Bogardus C, Lillioja S, Howard BV, Reaven G, Mott D. Relationships between insulin secretion, insulin action, and fasting plasma glucose concentration in nondiabetic and noninsulin-dependent diabetic subjects. J Clin Invest. 1984;74(4):1238–1246.

21. Porte D, Jr. Sympathetic regulation of insulin secretion. Its relation to diabetes mellitus. Arch Intern Med. 1969;123(3):252–260.

22. Gerich JE, Langlois M, Noacco C, Schneider V, Forsham PH. Adrenergic modulation of pancreatic glucagon secretion in man. J Clin Invest. 1974;53(5):1441–1446.

23. Clutter WE, Rizza RA, Gerich JE, Cryer PE. Regulation of glucose metabolism by sympathochromaffin catecholamines. Diabetes Metab Rev. 1988;4(1):1–15.

24. Sacca L, Vigorito C, Cicala M, Corso G, Sherwin RS. Role of gluconeogenesis in epinephrine-stimulated hepatic glucose production in humans. Am J Physiol. 1983;245(3):E294–302.

25. Silverberg AB, Shah SD, Haymond MW, Cryer PE. Norepinephrine: hormone and neurotransmitter in man. Am J Physiol. 1978;234(3):E252–256.

26. Sacca L, Morrone G, Cicala M, Corso G, Ungaro B. Influence of epinephrine, norepinephrine, and isoproterenol on glucose homeostasis in normal man. J Clin Endocrinol Metab. 1980;50(4):680–684.

27. Christensen NJ, Alberti KG, Brandsborg O. Plasma catecholamines and blood substrate concentrations: studies in insulin induced hypoglycaemia and after adrenaline infusions. Eur J Clin Invest. 1975;5(5):415–423.

28. Abramson EA, Arky RA. Role of beta-adrenergic receptors in counterregulation to insulin-induced hypoglycemia. Diabetes. 1968;17(3):141–146.

29. Rizza RA, Cryer PE, Haymond MW, Gerich JE. Adrenergic mechanisms for the effects of epinephrine on glucose production and clearance in man. J Clin Invest. 1980;65(3):682–689.

30. Exton JH, Robison GA, Sutherland EW, Park CR. Studies on the role of adenosine 3’,5’-monophosphate in the hepatic actions of glucagon and catecholamines. J Biol Chem. 1971;246(20):6166–6177.

31. Gerich J, Cryer P, Rizza R. Hormonal mechanisms in acute glucose counterregulation: the relative roles of glucagon, epinephrine, norepinephrine, growth hormone, and cortisol. Metabolism. 1980;29(11 Suppl 1):1164–1175.

32. Winkler H, Fischer-Colbrie R. The chromogranins A and B: the first 25 years and future perspectives. Neuroscience. 1992;49(3):497–528.

33. Bandyopadhyay GK, Mahata SK. Chromogranin A Regulation of Obesity and Peripheral Insulin Sensitivity. Front Endocrinol (Lausanne). 2017;8:20.

34. Muntjewerff EM, Dunkel G, Nicolasen MJT, Mahata SK, van den Bogaart G. Catestatin as a Target for Treatment of Inflammatory Diseases. Front Immunol. 2018;9:2199.

35. Mahata SK, Corti A. Chromogranin A and its fragments in cardiovascular, immunometabolic, and cancer regulation. Ann N Y Acad Sci. 2019;1455(1):34–58.

36. Taupenot L, Harper KL, O’Connor DT. The chromogranin-secretogranin family. New England Journal of Medicine. 2003;348(12):1134–1149.

37. Tatemoto K, Efendic S, Mutt V, Makk G, Feistner GJ, Barchas JD. Pancreastatin, a novel pancreatic peptide that inhibits insulin secretion. Nature. 1986;324(6096):476–478.

38. O’Connor DT, Cadman PE, Smiley C, et al. Pancreastatin: multiple actions on human intermediary metabolism in vivo, variation in disease, and naturally occurring functional genetic polymorphism. J Clin Endocrinol Metab. 2005;90(9):5414–5425.

39. Mahata SK, O’Connor DT, Mahata M, et al. Novel autocrine feedback control of catecholamine release. A discrete chromogranin A fragment is a noncompetitive nicotinic cholinergic antagonist. J Clin Invest. 1997;100(6):1623–1633.

40. Ying W, Mahata S, Bandyopadhyay GK, et al. Catestatin Inhibits Obesity-Induced Macrophage Infiltration and Inflammation in the Liver and Suppresses Hepatic Glucose Production, Leading to Improved Insulin Sensitivity. Diabetes. 2018;67(5):841–848.

41. Luketin M, Mizdrak M, Boric-Skaro D, et al. Plasma Catestatin Levels and Advanced Glycation End Products in Patients on Hemodialysis. Biomolecules. 2021;11(3).

42. Gayen JR, Saberi M, Schenk S, et al. A novel pathway of insulin sensitivity in chromogranin a null mice: A crucial role for pancreastatin in glucose homeostasis. J Biol Chem. 2009;284:28498–28509.

43. Bandyopadhyay GK, Lu M, Avolio E, et al. Pancreastatin-dependent inflammatory signaling mediates obesity-induced insulin resistance. Diabetes. 2015;64(1):104–116.

44. Pereda D, Pardo MR, Morales Y, Dominguez N, Arnau MR, Borges R. Mice lacking chromogranins exhibit increased aggressive and depression-like behaviour. Behavioural brain research. 2015;278:98–106.

45. Dasgupta A, Bandyopadhyay GK, Ray I, et al. Catestatin improves insulin sensitivity by attenuating endoplasmic reticulum stress: In vivo and in silico validation. Comput Struct Biotechnol J. 2020;18:464–481.

46. Mahata SK, Mahata M, Wakade AR, O’Connor DT. Primary structure and function of the catecholamine release inhibitory peptide catestatin (chromogranin A344-364): Identification of amino acid residues crucial for activity. Mol Endocrinol. 2000;14(10):1525–1535.

47. Mahata SK, Mahapatra NR, Mahata M, et al. Catecholamine secretory vesicle stimulus-transcription coupling in vivo. Demonstration by a novel transgenic promoter/photoprotein reporter and inhibition of secretion and transcription by the chromogranin A fragment catestatin. J Biol Chem. 2003;278:32058–32067.

48. Wen G, Mahata SK, Cadman P, et al. Both rare and common polymorphisms contribute functional variation at CHGA, a regulator of catecholamine physiology. Am J Hum Genet. 2004;74(2):197–207.

49. Mahata SK, Mahata M, Fung MM, O’Connor DT. Catestatin: a multifunctional peptide from chromogranin A. Regul Pept. 2010;162(1-3):33–43.

50. DeFronzo RA, Jacot E, Jequier E, Maeder E, Wahren J, Felber JP. The effect of insulin on the disposal of intravenous glucose. Results from indirect calorimetry and hepatic and femoral venous catheterization. Diabetes. 1981;30(12):1000–1007.

51. Jue T, Rothman DL, Tavitian BA, Shulman RG. Natural-abundance 13C NMR study of glycogen repletion in human liver and muscle. Proc Natl Acad Sci U S A. 1989;86(5):1439–1442.

52. Marchand I, Chorneyko K, Tarnopolsky M, et al. Quantification of subcellular glycogen in resting human muscle: granule size, number, and location. J Appl Physiol (1985). 2002;93(5):1598–1607.

53. Hokken R, Laugesen S, Aagaard P, et al. Subcellular localization- and fibre type-dependent utilization of muscle glycogen during heavy resistance exercise in elite power and Olympic weightlifters. Acta Physiol (Oxf). 2021;231(2):e13561.

54. Rui L. Energy metabolism in the liver. Compr Physiol. 2014;4(1):177–197.

55. Ciudad CJ, Carabaza A, Guinovart JJ. Glucose 6-phosphate plays a central role in the activation of glycogen synthase by glucose in hepatocytes. Biochem Biophys Res Commun. 1986;141(3):1195–1200.

56. Carabaza A, Ciudad CJ, Baque S, Guinovart JJ. Glucose has to be phosphorylated to activate glycogen synthase, but not to inactivate glycogen phosphorylase in hepatocytes. FEBS Lett. 1992;296(2):211–214.

57. Villar-Palasi C, Guinovart JJ. The role of glucose 6-phosphate in the control of glycogen synthase. FASEB J. 1997;11(7):544–558.

58. Ferrer JC, Favre C, Gomis RR, et al. Control of glycogen deposition. FEBS Lett. 2003;546(1):127–132.

59. Aiston S, Green A, Mukhtar M, Agius L. Glucose 6-phosphate causes translocation of phosphorylase in hepatocytes and inactivates the enzyme synergistically with glucose. Biochem J. 2004;377(Pt 1):195–204.

60. Villar-Palasi C. Substrate specific activation by glucose 6-phosphate of the dephosphorylation of muscle glycogen synthase. Biochim Biophys Acta. 1991;1095(3):261–267.

61. Bouskila M, Hunter RW, Ibrahim AF, et al. Allosteric regulation of glycogen synthase controls glycogen synthesis in muscle. Cell Metab. 2010;12(5):456–466.

62. Prats C, Cadefau JA, Cusso R, et al. Phosphorylation-dependent translocation of glycogen synthase to a novel structure during glycogen resynthesis. J Biol Chem. 2005;280(24):23165–23172.

63. Poulter L, Ang SG, Gibson BW, et al. Analysis of the in vivo phosphorylation state of rabbit skeletal muscle glycogen synthase by fast-atom-bombardment mass spectrometry. Eur J Biochem. 1988;175(3):497–510.

64. Roach PJ. Control of glycogen synthase by hierarchal protein phosphorylation. FASEB J. 1990;4(12):2961–2968.

65. Jope RS, Johnson GV. The glamour and gloom of glycogen synthase kinase-3. Trends Biochem Sci. 2004;29(2):95–102.

66. Hughes K, Nikolakaki E, Plyte SE, Totty NF, Woodgett JR. Modulation of the glycogen synthase kinase-3 family by tyrosine phosphorylation. EMBO J. 1993;12(2):803–808.

67. Cross DA, Alessi DR, Cohen P, Andjelkovich M, Hemmings BA. Inhibition of glycogen synthase kinase-3 by insulin mediated by protein kinase B. Nature. 1995;378(6559):785–789.

68. Jensen J, Brennesvik EO, Lai YC, Shepherd PR. GSK-3beta regulation in skeletal muscles by adrenaline and insulin: evidence that PKA and PKB regulate different pools of GSK-3. Cell Signal. 2007;19(1):204–210.

69. Lawrence JC, Jr., Roach PJ. New insights into the role and mechanism of glycogen synthase activation by insulin. Diabetes. 1997;46(4):541–547.

70. Taniguchi CM, Emanuelli B, Kahn CR. Critical nodes in signalling pathways: insights into insulin action. Nat Rev Mol Cell Biol. 2006;7(2):85–96.

71. Storz P, Toker A. 3’-phosphoinositide-dependent kinase-1 (PDK-1) in PI 3-kinase signaling. Front Biosci. 2002;7:d886–902.

72. Gan X, Wang J, Su B, Wu D. Evidence for direct activation of mTORC2 kinase activity by phosphatidylinositol 3,4,5-trisphosphate. J Biol Chem. 2011;286(13):10998–11002.

73. Yang G, Murashige DS, Humphrey SJ, James DE. A Positive Feedback Loop between Akt and mTORC2 via SIN1 Phosphorylation. Cell Rep. 2015;12(6):937–943.

74. Copp J, Manning G, Hunter T. TORC-specific phosphorylation of mammalian target of rapamycin (mTOR): phospho-Ser2481 is a marker for intact mTOR signaling complex 2. Cancer Res. 2009;69(5):1821–1827.

75. Moore MC, Cherrington AD, Cline G, et al. Sources of carbon for hepatic glycogen synthesis in the conscious dog. J Clin Invest. 1991;88(2):578–587.

76. Agius L. Glucokinase and molecular aspects of liver glycogen metabolism. Biochem J. 2008;414(1):1–18.

77. Chu CA, Sindelar DK, Neal DW, Cherrington AD. Direct effects of catecholamines on hepatic glucose production in conscious dog are due to glycogenolysis. Am J Physiol. 1996;271(1 Pt 1):E127–137.

78. Syed NA, Khandelwal RL. Reciprocal regulation of glycogen phosphorylase and glycogen synthase by insulin involving phosphatidylinositol-3 kinase and protein phosphatase-1 in HepG2 cells. Mol Cell Biochem. 2000;211(1-2):123–136.

79. Dent P, Lavoinne A, Nakielny S, Caudwell FB, Watt P, Cohen P. The molecular mechanism by which insulin stimulates glycogen synthesis in mammalian skeletal muscle. Nature. 1990;348(6299):302–308.

80. Chen M, Halter JB, Porte D, Jr. Plasma catecholamines, dietary carbohydrate, and glucose intolerance: a comparison between young and old men. J Clin Endocrinol Metab. 1986;62(6):1193–1198.

81. Sarbassov DD, Guertin DA, Ali SM, Sabatini DM. Phosphorylation and regulation of Akt/PKB by the rictor-mTOR complex. Science. 2005;307(5712):1098–1101.

82. Manning BD, Toker A. AKT/PKB Signaling: Navigating the Network. Cell. 2017;169(3):381–405.

83. Gayen JR, Gu Y, O’Connor DT, Mahata SK. Global disturbances in autonomic function yield cardiovascular instability and hypertension in the chromogranin A null mouse. Endocrinology. 2009;150(11):5027–5035.

84. Bandyopadhyay GK, Vu CU, Gentile S, et al. Catestatin (chromogranin A(352-372)) and novel effects on mobilization of fat from adipose tissue through regulation of adrenergic and leptin signaling. J Biol Chem. 2012;287(27):23141–23151.

85. Rabbi MF, Munyaka PM, Eissa N, Metz-Boutigue MH, Khafipour E, Ghia JE. Human Catestatin Alters Gut Microbiota Composition in Mice. Front Microbiol. 2016;7:2151.

86. Rabbi MF, Eissa N, Munyaka PM, et al. Reactivation of Intestinal Inflammation Is Suppressed by Catestatin in a Murine Model of Colitis via M1 Macrophages and Not the Gut Microbiota. Front Immunol. 2017;8:985.

87. Muntjewerff EM, Tang K, Lutter L, et al. Chromogranin A regulates gut permeability via the antagonistic actions of its proteolytic peptides. Acta Physiol (Oxf). 2021;232(2):e13655.

88. Ying W, Tang K, Avolio E, et al. Immunosuppression of Macrophages Underlies the Cardioprotective Effects of CST (Catestatin). Hypertension. 2021;77(5):1670–1682.

89. Riopel M, Seo JB, Bandyopadhyay GK, et al. Chronic fractalkine administration improves glucose tolerance and pancreatic endocrine function. J Clin Invest. 2018;128(4):1458–1470.

90. Carroll NV, Longley RW, Roe JH. The determination of glycogen in liver and muscle by use of anthrone reagent. J Biol Chem. 1956;220(2):583–593.

91. Ciraolo E, Gulluni F, Hirsch E. Methods to measure the enzymatic activity of PI3Ks. Methods Enzymol. 2014;543:115–140.

