## Supplemental Figures_BioRxiv_110421 for "Catestatin induces glycogenesis by stimulating phosphoinositide 3-kinase-AKT pathway"

[1]VA San Diego Healthcare System, 3350 La Jolla Village Drive, San Diego, CA, USA; [2]Department of Medicine, University of California San Diego, La Jolla, CA, USA; [3]Department of Tumor Immunology, Radboud Institute for Molecular Life Sciences, Radboud University Medical Center, Nijmegen, the Netherlands; [4]Department of Molecular Immunology and Microbiology, Groningen Biomolecular Sciences and Biotechnology Institute, University of Groningen, Groningen, the Netherlands.

*Contributed equally to this work.

**Keywords:** Catestatin; Phospho-AKT, Hyperglycemia; Glycogen; Glucose-6-phosphate; Glycogen synthase

**Short title:** Catestatin diverts gluconeogenic substrates to the glycogenic pathway

**¶Correspondence should be addressed to**:

Sushil K. Mahata, Ph.D.
Metabolic Physiology & Ultrastructural Biology Laboratory
Department of Medicine
University of California San Diego
9500 Gilman Drive
La Jolla, CA 92093-0732

**Figure S1:** TEM micrographs showing glycogen granules in liver after treatments with saline or CST in (A&B) Fed and (C&D) fasted NCD-WT mice. (E) Morphometric analyses of the glycogen granules. (F) Dose-dependent increase in glycogen synthesis in fasted NCD-WT mice by CST. (G) Half-life of plasma CST in Fed NCD-WT mice.

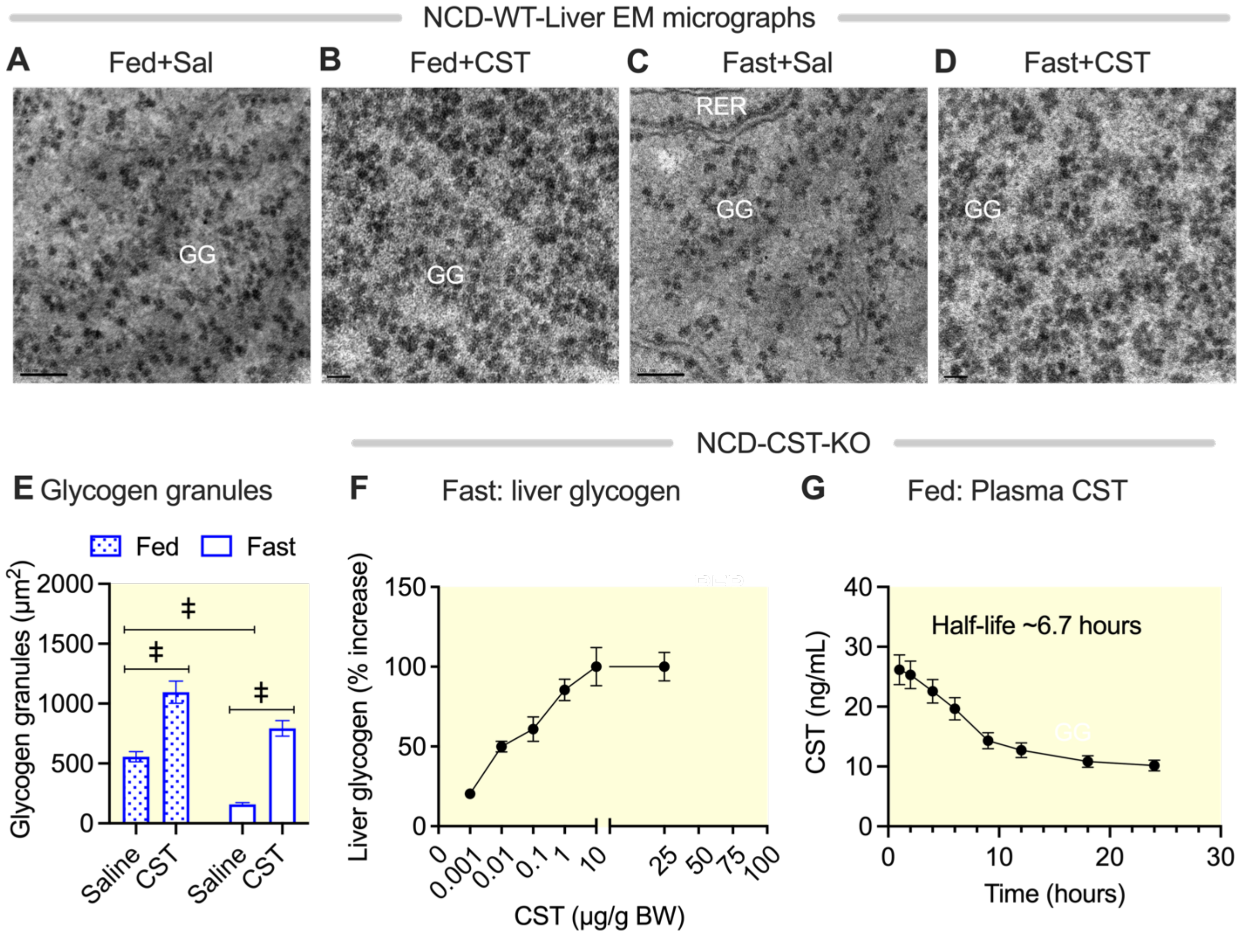
NCD-WT-Liver EM micrographs
A
Fed+Sal
B
Fed+CST
C
Fast+Sal
D
Fast+CST
GG
RER
NCD-CST-KO
E Glycogen granules
Fed
Fast
Glycogen granules (μm²)
2000
1500
1000
500
0
‡
Saline
CST
F
Fast: liver glycogen
Liver glycogen (% increase)
150
100
50
0
0.001
0.01
0.1
1
10
25
50
75
100
CST (μg/g BW)
G
Fed: Plasma CST
Half-life ~6.7 hours
CST (ng/mL)
40
30
20
10
0
Time (hours)

Figure S1

**Figure S2:** TEM micrographs showing glycogen granules in gastrocnemius muscles after treatments with saline or CST: (A&B) sub-sarcolemmal region; (C&D) myofibrillar region. TEM micrographs showing lipid droplets in DIO-WT liver after treatments with saline or CST: (E) saline; (F) CST.

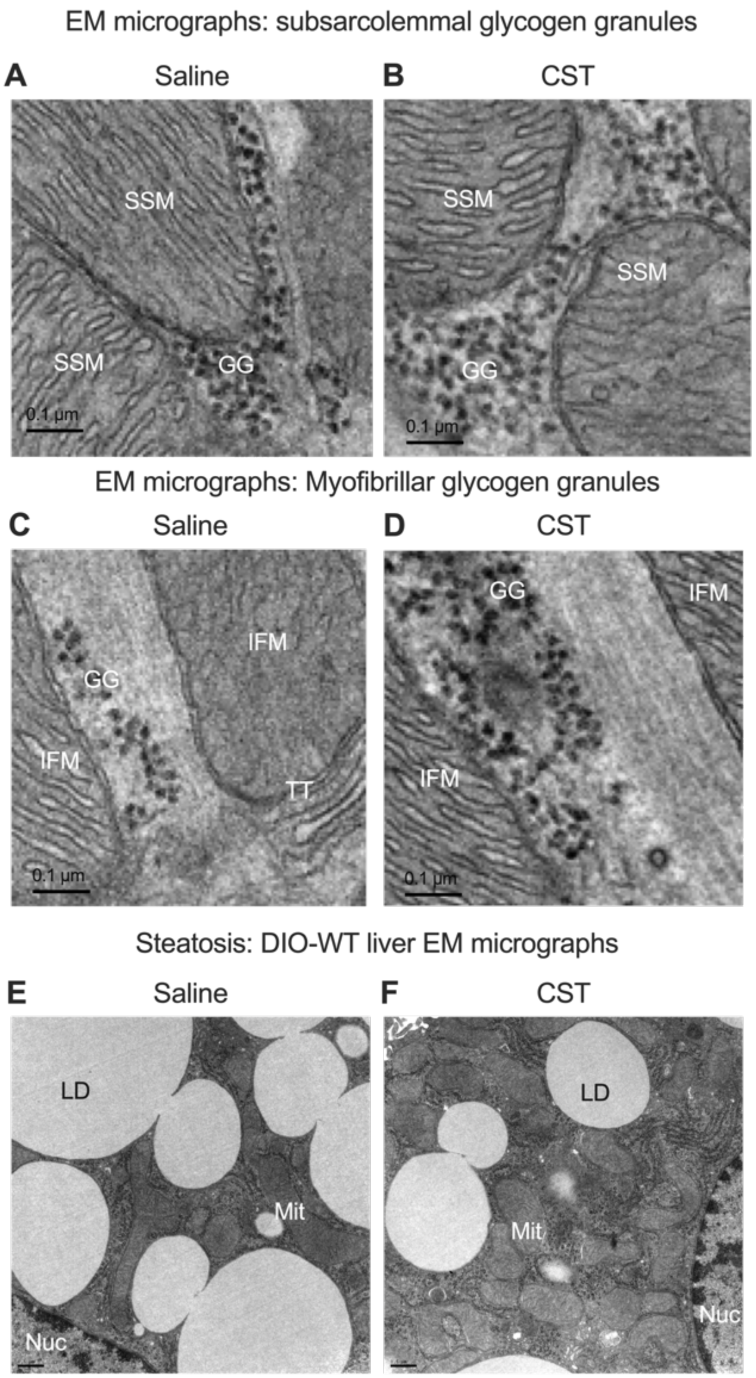
EM micrographs: subsarcolemmal glycogen granules
A
Saline
B
CST
SSM
SSM
GG
0.1 μm
SSM
SSM
GG
0.1 μm
EM micrographs: Myofibrillar glycogen granules
C
Saline
D
CST
IFM
GG
IFM
TT
0.1 μm
GG
IFM
IFM
0.1 μm
Steatosis: DIO-WT liver EM micrographs
E
Saline
F
CST
LD
Mit
Nuc
LD
Mit
Nuc

Figure S2

**Figure S3:** Plasma levels of counter-regulatory hormones (NE, EPI, and glucagon) in the fed and fasted DIO-WT mice after treatments with saline or CST.

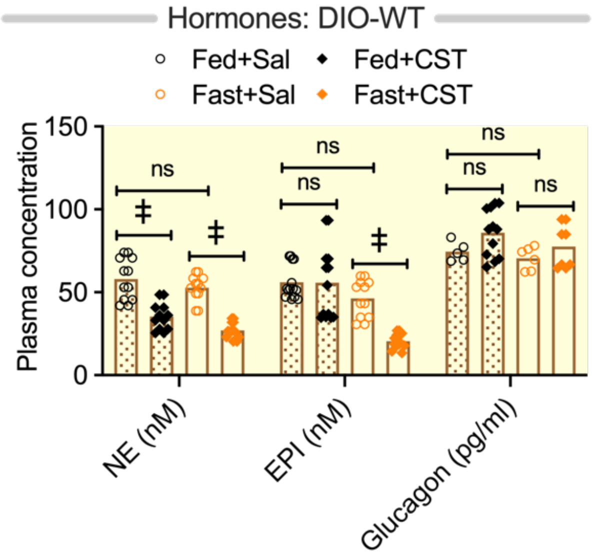

Hormones: DIO-WT
Fed+Sal
Fed+CST
Fast+Sal
Fast+CST
Plasma concentration
150
100
50
0
ns
‡
ns
ns
ns
ns
ns
NE (nM)
EPI (nM)
Glucagon (pg/ml)


Figure S3

**Figure S4:** Effects of oral CST treatment in DIO-WT MICE on (A) body weight and hepatic expression of gluconeogenic genes: (B) *G6pc* (B) and (C) *Pck-1*.

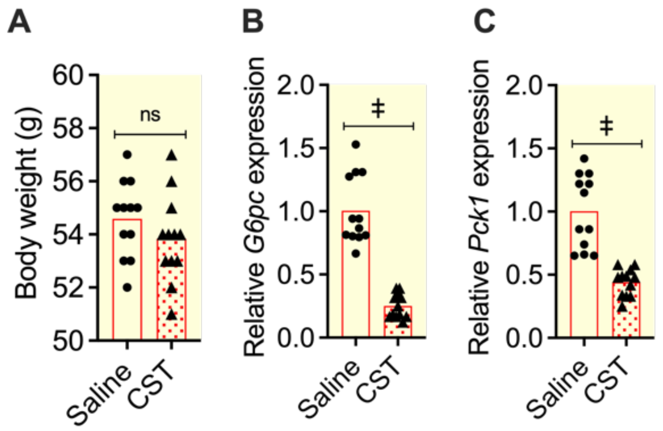
A
B
C
Body weight (g)
ns
Saline
CST
Relative G6pc expression
‡
Relative Pck1 expression


Figure S4
